# Information-Content-Informed Kendall-tau Correlation Methodology: Interpreting Missing Values in Metabolomics as Potentially Useful Information

**DOI:** 10.1101/2022.02.24.481854

**Authors:** Robert M Flight, Praneeth S Bhatt, Hunter NB Moseley

**Affiliations:** Markey Cancer Center, University of Kentucky, Lexington, KY 40536, USA; Department of Molecular & Cellular Biochemistry, University of Kentucky, Lexington, KY 40536, USA; Superfund Research Center, University of Kentucky, Lexington, KY 40536, USA; Department of Electrical and Computer Engineering, University of Kentucky, Lexington, KY 40506; Institute for Biomedical Informatics, University of Kentucky, Lexington, KY 40536, USA; Department of Toxicology and Cancer Biology, University of Kentucky, Lexington, KY 40536, USA

**Keywords:** Metabolomics, Correlation, Missingness, Left-Censored

## Abstract

**Background:** Almost all correlation measures currently available are unable to directly handle missing values. Typically, missing values are either ignored completely by removing them or are imputed and used in the calculation of the correlation coefficient. In either case, the correlation value will be impacted based on a perspective that the missing data represents no useful information. However, missing values occur in real data sets for a variety of reasons. In metabolomics data sets, a major reason for missing values is that a specific measurable phenomenon falls below the detection limits of the analytical instrumentation (left-censored values). These missing data are not missing at random, but represent potentially useful information by virtue of their “missingness” at one end of the data distribution.

**Methods:** To include this information due to left-censorsed missingness, we propose the information-content-informed Kendall-tau (ICI-Kt) methodology. We developed a statistical test and then show that most missing values in metabolomics datasets are the result of left-censorship. Next, we show how left-censored missing values can be included within the definition of the Kendall-tau correlation coefficient, and how that inclusion leads to an interpretation of information being added to the correlation. We also implement calculations for additional measures of theoretical maxima and pairwise completeness that add further layers of information interpretation in the methodology.

**Results:** Using both simulated and over 700 experimental data sets from The Metabolomics Workbench, we demonstrate that the ICI-Kt methodology allows for the inclusion of left-censored missing data values as interpretable information, enabling both improved determination of outlier samples and improved feature-feature network construction.

**Conclusions:** We provide explicitly parallel implementations in both R and Python that allow fast calculations of all the variables used when applying the ICI-Kt methodology on large numbers of samples. The ICI-Kt methods are available as an R package and Python module on GitHub at https://github.com/moseleyBioinformaticsLab/ICIKendallTau and https://github.com/moseleyBioinformaticsLab/icikt, respectively.

## 1. Introduction

Correlation as a measure of the relatedness or similarity of two or more sets of data has a long history, with the mathematical technique being used (and abused) in various scientific fields since its introduction [1,2]. More recently, correlation calculations have become a cornerstone statistical method in the analysis and integration of varied omics’ datasets, especially the big five omics: genomics, transcriptomics, proteomics, metabolomics, and epigenomics [3]. Correlation between biomolecular features (nucleotide variants, RNA transcripts, proteins, metabolites, DNA methylation, etc.) may be used to evaluate the relationship strength between pairs of the features as well as to detect and derive correlative structures between groups of features [4]. Moreover, feature-feature correlations can be used to evaluate a dataset based on expected biochemical correlations, for example higher feature-feature correlations within lipid categories versus between lipid categories [5]. Correlation is a foundational method for generating biomolecular feature-feature interaction networks, like those provided by STRING [6], Genemania [7], and WCGNA [8]. Feature-feature correlation may also be used to inform which features are used for imputation of missing values [9].

Often, the first step in omics’ level analyses is to examine the sample-sample (dis)similarities in various ways using exploratory data analysis or EDA. This can include the examination of decomposition by principal components analysis (PCA), sample-sample pairwise distances, or sample-sample pairwise correlations to highlight biological and batch groups [10–12], double check the appropriateness of planned analyses [13], and check if any samples should be removed prior to statistical analysis (outlier detection and removal) [14]. Outlier detection, in particular, is often required for successful omics’ data analysis, as any misstep during the experimentation, sample collection, sample preparation, or analytical measurement of individual samples can inject high error and/or variance into the resulting data set [10–12,14,15].

All analytical methods, and in particular the analytical methods used in omics’ where many analytes are being measured simultaneously, suffer from missing measurements. Some analytes will be missing at random because of spurious issues with either the instrument, the particular sample, or sample preparation, but a larger number of missing measurements are left-censored due to analytes being below the effective detection limit of the instrument and the given specific sample preparation procedures utilized, as shown in Figure 1. Some analytical instruments are purposely designed to floor measurements when they occur below a certain signal to noise ratio threshold. Also, imputation of missing measurements in omics’ samples is an active area of research, which we will not comprehensively cover here beyond to say that it is worthwhile and very necessary in many instances. Imputation methods rely on very similar analytical detection limits between analytical samples. When this condition does not hold, imputation methods have reduced performance and lower interpretive value. For analytical techniques requiring complex sample handling and detection, the variability in the analytical detection level can be quite high. However, when it comes to calculating correlation, there are very few methods that explicitly account for left-censored missing data that we know of. In many cases, missing values are either ignored or imputed to zero (or another value) and then included in the correlation calculation. The two most common approaches for ignoring (i.e. dropping) values is to only use those measurements that are common across all samples (complete) or that are common between two samples being compared (pairwise complete). Both dropping or imputing missing values are likely to cause the calculated sample-sample correlation values to deviate from the real sample-sample correlation values, especially with respect to specific data interpretation perspectives.

**Figure 1.**
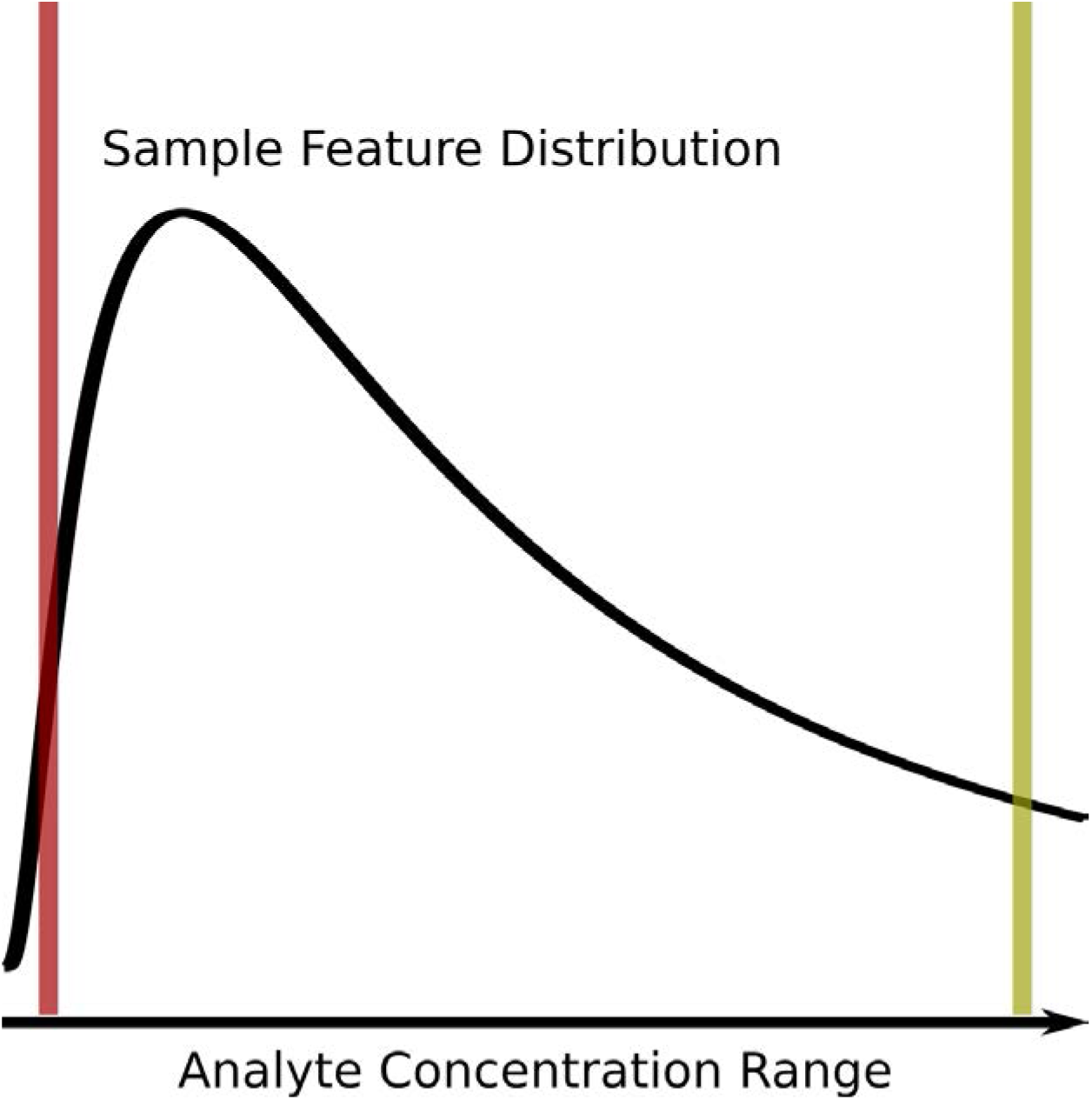
Graphical description of the left-censored data problem. An example density plot of the analyte concentrations for a single experimental sample is shown as a solid black curve. The true analyte concentration range covers the full range of the density distribution, with the minimum on the left (red vertical line), and the maximum on the right (yellow vertical line). Below certain concentrations, shown by the red line, the instrument returns either missing values (NA), zeros, or some other floored values, resulting in a left-censored distribution. Above certain concentrations, highlighted by the yellow line, typically the values returned will be identical (or flagged depending on the instrument). Which analytes will have concentrations below the red detection limit line may vary from sample to sample due to the overall sample composition, as well as biological variance.

Assuming that a majority of missing values are not missing at random, but rather result from left-censored distributions due to the analyte being below the effective detection limit (see Figure 1), we propose that these missing values do in fact encode useful information that can be incorporated into correlation calculations.

To create a correlation measure that is capable of working with missing values, we would not be interested in creating a completely new correlation metric from scratch, but modifying an existing one. Of the three commonly used correlation measures, Pearson, Spearman, and Kendall-tau, Spearman and Kendall-tau seem most appropriate for modification as they solely use ranks in the calculation of their coefficients. Modifying Pearson would either involve imputing new values, or finding a way to calculate the covariances **with** missingness included. While Spearman uses ranks, many of the modifications for handling identical ranks and ties do not seem amenable to working with missing values. In contrast, Kendall-tau’s use of concordant and discordant pair counts seems most amenable to the creation of new definitions that incorporate missingness while still working within the original definition of the correlation coefficient, as shown in the Implementation section below.

In this work, we propose new definitions of concordant and discordant rank pairs that include missing values, as well as methods for incorporating missing values into the number of tied values for the equivalent of the modified Kendall-tau coefficient, the information-content-informed Kendall-tau (ICI-Kt) method. The implementation of the basic calculation of ICI-Kt involves the replacement of missing values with a value lower than the observed values (technically simple imputation), with subsequent calculation of the Kendall *τ*_*b*_ statistic, as a majority of missing values are the result of left-censorship, they still provide an interpretation from an information-content perspective, which we demonstrate with the equations below. We also developed a binomial statistical test for determining if the cause for missingness is likely left-censorship. With this statistical test and experimental datasets from The Metabolomics Workbench (MW) [16], we demonstrate that left-censorship is the cause of many missing values across a large number of metabolomics datasets. We examine the effect of missing values on various collections of simulated and real data-sets, comparing the ICI-Kt methodology with other simpler methods of handling the missing values, namely removing them or imputing them to zero. Given the detrimental effects of including outlier samples in differential analyses, we also evaluate the ability of the ICI-Kt methodology to capture sample-sample pairwise similarities and the determination of outlier samples prior to differential analyses. We were also curious about the utility of the ICI-Kt methodology in creating metabolomics feature-feature networks with large amounts of missing values, so we evaluated the partitioning of networks by Reactome pathways [17] after network creation using different correlation measures.

All of the code used for this manuscript is available on zenodo [18].

## 2. Materials and Methods

### 2.1 Additional Definitions of Concordant and Discordant Pairs to Include Missingness

In the simplest form, the Kendall-tau (*τ*_*a*_) correlation can be defined as:

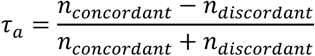

where *n*_*concordant*_ is the the number of concordant pairs and *n*_*discordant*_ is the number of discordant pairs. In this case, a pair are any two x-y points, *x*_*i*_, *y*_*i*_ and *x*_*j*_, *y*_*j*_, with *i* ≠ *j*, composed from two joint random variables X and Y, where *x*_*i*_ represents the *ith value* in X and *y*_*i*_ represents the *ith value* in Y. In a metabolomics context, X and Y can represent metabolite feature vectors for two experimental samples or two specific metabolite features across a set of samples.

A concordant pair has the following classical definition:

_•_ *x*_*j*_ and *y*_*i*_ > *y*_*j*_
_•_ *x*_*i*_ < *x*_*j*_ and *y*_*i*_ < *y*_*j*_

A discordant pair has the following classical definition [19]:

_•_ *x*_*j*_ and *y*_*i*_ < *y*_*j*_
_•_ *x*_*i*_ < *x*_*j*_ and *y*_*i*_ > *y*_*j*_

We can expand the concordant and discordant pair definitions to include missing values (e.g. NA in R). Here ! *x* indicates *x* = NA and ! *y* indicates *y* = NA, and & is synonymous with “and”. The information-content-informed concordant pair definitions are then:

_•_ *x*_*j*_ and *y*_*i*_ > *y*_*j*_
_•_ *x*_*i*_ < *x*_*j*_ and *y*_*i*_ < *y*_*j*_
_•_ *x*_*i*_ > *x*_*j*_ and *y*_*i*_&! *y*_*j*_
_•_ *x*_*i*_ < *x*_*j*_ and ! *y*_*i*_&*y*_*j*_
_•_ *x*_*i*_&! *x*_*j*_ and *y*_*i*_ > *y*_*j*_
_•_ ! *x*_*i*_&*x*_*j*_ and *y*_*i*_ < *y*_*j*_
_•_ *x*_*i*_&! *x*_*j*_ and *y*_*i*_&! *y*_*j*_
_•_ ! *x*_*i*_&*x*_*j*_ and ! *y*_*i*_&*y*_*j*_

The information content informed discordant pair definitions are then:

_•_*x*_*i*_ > *x*_*j*_ and *y*_*i*_ < *y*_*j*_
_•_ *x*_*i*_ < *x*_*j*_ and *y*_*i*_ > *y*_*j*_
_•_ *x*_*i*_ > *x*_*j*_ and ! *y*_*i*_&*y*_*j*_
_•_ *x*_*i*_ < *x*_*j*_ and *y*_*i*_&! *y*_*j*_
_•_ *x*_*i*_&! *x*_*j*_ and *y*_*i*_ < *y*_*j*_
_•_ ! *x*_*i*_&*x*_*j*_ and *y*_*i*_ > *y*_*j*_
_•_ *x*_*i*_&! *x*_*j*_ and ! *y*_*i*_&*y*_*j*_
_•_ ! *x*_*i*_&*x*_*j*_ and *y*_*i*_&! *y*_*j*_

These additional definitions make it possible to interpret a Kendall-tau correlation from the perspective of **missing values as additional information**, i.e., information-content-informed Kendall-tau (ICI-Kt) methodology.

### 2.2 Considering Ties

Tied values do not contribute to either of the concordant or discordant pair counts, and the original Kendall-tau formula for the *τ*_*a*_ statistic does not consider the presence of tied values. However, the related *τ*_*b*_ statistic does handle the presence of tied values by adding the tied *x* and *y* values to the denominator, and in our special case of missing data, we can add the ties that result from (*x*_*i*_ = NA, *x*_*j*_ = NA) and (*y*_*i*_ = NA, *y*_*j*_ = NA) to *n*_*xtie*_ and *n*_*ytie*_ [20,21] used in the following equation for *τ*_*b*_:

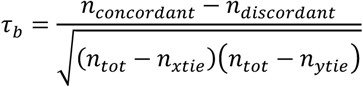

where *n*_*tot*_ is the total number of pairs, *n*_*xtie*_ are the number of tied values in X, and *n*_*ytie*_ are the number of paired values in Y. We can also consider commonly missing values in X and Y specially as well. In the first instance, we remove those x-y points where **both values** are missing. We refer to this case as the *local* ICI-Kt correlation. It is most appropriate for the comparison of only two experimental samples, where we are concerned with what values are present in the two experimental samples, with the odd case of missingness.

The other case, where we leave ties resulting from points with missing X and Y, we refer to as the *global* ICI-Kt correlation. In this case, every single correlation over multiple comparisons with the same set of metabolite features will consider the same number of pair comparisons. This is useful when analyzing and interpreting correlations from a large number of experimental samples, not just two samples.

### 2.3 Theoretical Maxima

The *global* case also provides an interesting property, whereby we can calculate the theoretical maximum correlation that would be possible to observe given the lowest number of shared missing values. This value can be useful to scale the rest of the observed correlation values across many sample-sample correlations, providing a value that scales an entire dataset appropriately. For any pairwise comparison of two vectors (from experimental samples for example), we can calculate the maximum possible Kendall-tau for that comparison by defining the maximum number of concordant pairs as:

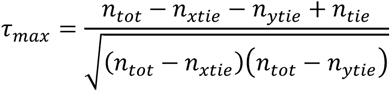

where *n*_*tie*_ is the number of commonly tied values in both X and Y. Calculating a set of *τ*_*max*_ values between all experimental samples, we can take the maximum of the values, and use it to scale **all** of the obtained Kendall-tau values equally 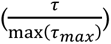.

### 2.4 Completeness

As an addition to the correlation value, we also calculate the *completeness* between any two samples. We first measure the number of entries missing in either of the samples being compared, and subtract that from the total number of features in the samples. This defines how many features are potentially *complete* between the two samples. This number, over the total number of features defines the *completeness* fraction.

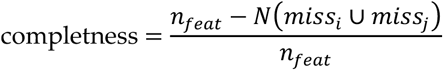

where for any two samples *i* and *j*, *n*_*feat*_ is the total number of features or entries, and *miss*_*i*_ ∪ *miss*_*j*_ are the metabolite features missing in either sample *i* or *j*, with *N* being the total number of missing entries in either sample *i* or *j*.

### 2.5 Implementation Details

We produced an initial reference implementation in base R [22], where the various concordant and discordant pair definitions were written as simple logical tests to allow further exploration and validation of faster implementations. During exploration and validation of an early implementation, we discovered that an equivalent calculation was to replace the missing values with a value smaller than all of the values in the two sets being compared. This simplification does not change the interpretation of the effect of left-censored missing values, but it does allow for the direct use of the very fast mergesort based algorithm for calculating *τ*_*b*_ [23].

We re-implemented the mergesort implementation from the SciPy kendalltau code [24] in both R (via Rcpp) and Python (via cython) to enable fast, easy parallel computations in both languages (using furrr and multiprocessing, respectively), as well as inclusion of the calculation of *tau*_*max*_, which is derived from the same values needed for the calculation of *τ*_*b*_ (see above). The version of the ICIKendallTau R package used in this manuscript is available on zenodo [25]. In addition to use as an imported Python module, the Python icikt module provides a command line interface (CLI) for the ICI-Kt methodology.

### 2.6 Simulated Data Sets

Simulated feature vectors (analytical samples) are generated by drawing 1000 random values from a log-normal distribution with a mean of 1 and standard deviation (sd) of 0.5 and sorting them in ascending order to create a pair of samples with perfectly positive (1) or negative (-1) correlation values. Random variance is added to one of the two samples by drawing values from a uniform distribution over -0.5 to 0.5, and adding the values to the original sample, and sorting them again to maintain a correlation of 1 or -1 for Kendall-tau correlation. A sample with a small percentage (0.5%) outlier points at one end of the distribution is created by sampling from a uniform distribution over the range -0.5 to 0.5, and then a log-normal distribution with a mean of 1.2 and standard deviation of 0.1 for the number of desired outlier points, and adding the log-normal values to the uniform values for a combined source of random variance that is added to the original sample values. The negative analytical sample has values sorted in descending order. Missing value indices are generated by randomly sampling up to 499 of the lowest values in each sample. For the negative sample, the indices are also subtracted from 1000 to cause them to be at the lower end of the feature distribution. Finally, missing indices were only inserted into one of the two samples being compared before calculating the correlation. The missing indices are replaced with NA, and then correlations between the analytical samples are calculated.

Another, more realistic, simulated data set is generated by drawing 1000 random values from a log-normal distribution, and adding noise from a normal distribution with a mean of zero and sd of 0.2 to create two statistical samples. Missing values are created in these statistical samples via two methods: 1) by creating intensity cutoffs from 0 to 1.5 in 0.1 increments, values below the cutoff set to missing or zero depending on the calculation; 2) randomly sampling locations in the two-sample matrix ranging from zero to 300 in increments of 50 and setting the indices to missing or zero.

### 2.7 Metabolomics Datasets from Metabolomics Workbench

A set of 6105 analysis datasets from The Metabolomics Workbench (MW) were downloaded on 2025-11-12 using the mwtab python package [26], and repaired to fix various issues. For a subset that had metabolite feature abundances outside the mwtab json file, the files were downloaded on 2025-11-13. The various pieces of each dataset were parsed and transformed into R appropriate structures, mainly data frames of various types for metadata, and matrices of abundances (see Data Processing). Subject sample factors (SSF) were transformed so that each sample has a combination of various factors to describe the unique groups of samples. For example, if a dataset included samples with one or more genotypes (Knockout, FLOX) and taken from different segments of the intestines, the final factor for each sample is the combination of genotype + intestinal segment.

For inclusion in this work, an analysis dataset had to meet these criteria:

- ≥ 100 metabolites;
- one SSF grouping with ≥ 5 samples, and ≥ 2 SSF groupings after removing samples that **may** be pooled, quality-control or blanks;
- a maximum metabolite feature abundance ≥ 20 to exclude log-transformed values and low dynamic range datasets;
- the ability to calculate a correlation between the median rank of a metabolite feature and the number of samples the metabolite was missing in within a factor.

Of the 6105 datasets initially downloaded, 711 were kept for further analysis.

### 2.8 Number of Missing Values and Median Rank

For each dataset, the samples were split by SSF (see previous Methods). For each metabolite feature, the rank of the feature was calculated for each sample where the feature was present, followed by the features median rank across samples, as well as the number of samples the feature was missing from. Grouping the features by the number of missing values, we calculate the median of median ranks, as well as the minimum of median ranks, for the visualization and correlation of the relationship of rank with missing values.

### 2.9 Binomial Test for Left Censorship

For each dataset, the samples were first split by SSF. In each sample, the median abundance of features present in the sample are calculated. For any feature that is missing in any sample, the values in the present samples are compared to the median value of their corresponding sample. If the value is less than or equal to the median value in the sample, that is counted as a **success** in a binomial test, otherwise it is counted as a **failure**. The number of successes and failures are aggregated across the SSF splits for calculation in a binomial test, with the null hypothesis assuming a ratio of 0.5.

### 2.10 Correlation Methods

For each dataset, we calculated correlations using a variety of methods. Across the various datasets, either zeros or empty strings (generally resulting in NA values when read into R) are used to represent missingness. To start, we replaced all missing values with NA, and then either left them as NA or set them to zero for the various methods used. ICI-Kt with NA (icikt); and then scaled (multiplied) by the completeness metric (icikt_complete). Kendall-tau, with NA, and then using pairwise-complete-observations (kt_base). Pearson, with NA’s replaced by zeros, using pairwise-complete-observations (pearson_base). Pearson, with NA, using pairwise-complete-observations (pearson_base_nozero). Pearson, with a *log*(*x* + 1) transform applied, using pairwise-complete-observations (pearson_log1p). Pearson, with a *log*(*x*) transform, and then setting infinite values to NA values, using pairwise-complete-observations (pearson_log).

### 2.11 Outlier Detection

For outlier detection, median sample-sample correlations within the unique SSF (genotype, condition, and their combinations) is calculated, and *log*(1 − *cor*_*median*_) calculated to transform it into a score. Then outliers are determined using grDevices::box-plot.stats, which by default are at 1.5X the whiskers in a box-and-whisker plot. As we are interested in only those correlations at the *low* end of correlation (becoming the high end after the subtraction and log-transform), we restrict to only those entries at the *high* end of the score distribution (using visualizationQualityControl::determine_outliers [27]). This is equivalent to using the correlation component of the score described by Gierliński et al [14] and setting the other component weights to zero.

After outlier detection, the ANOVA statistics are calculated across SSF using limma (v 3.62.2) [28], and the fraction of metabolites with an adjusted p-value ≤ 0.05 recorded (Benjamini-Hochberg adjustment [29]). The correlation methods are compared by their fractions of significant metabolites, using a paired t-test, and adjusting p-values across method comparisons using the Bonferroni adjustment.

### 2.12 Feature Annotations

Predicted Reactome pathway annotations for analyzable MW datasets were parsed from the previous work by Huckvale et al. [17,30]. Given the hierarchical nature of Reactome pathways, and the use of the pathways for partitioning feature-feature networks, we aggregated the Reactome pathways into larger grouped pathway sets with less feature overlap. First, we aggregated predicted pathway annotations to the same pathway identifier across species. Second, for each pathway annotation with ≥ 20 and ≤ 500 features, we calculated the combined overlap metric between all pathways (categoryCompare2 v 0.200.4 [31,32], values range between 0 and 1). Treating the overlap metric between pathways as weighted edges of a graph, we removed the edges with a value ≤ 0.6, and then did community detection using the walktrap clustering method in the igraph package (v 2.1.4)[33–36]. Each community of Reactome pathways identified by the walktrap clustering were then aggregated to a new grouped pathway annotation. These grouped pathway annotations were then used for the network partitioning calculations.

### 2.13 Feature-Feature Networks and Partitioning

Each MW dataset that was used for outlier detection (711) was also checked against the list of datasets used in the prediction of pathways and enrichment by Huckvale et al. (resulting in 137 datasets) [30]. The various correlation measures were calculated between all features (see Correlation Methods). For any given correlation method, we generate the feature-feature network for that dataset-correlation method combination. The dataset correlations were transformed to partial correlations. From the distribution of partial correlation values, we consider the fraction of values that make up the 2.5 % of the tail values (for a total of 5%) as the **significant** partial correlations that can be used as actual edges in the network. The network is then trimmed to only the edges that have a positive weight.

For each feature annotation (see Feature Annotations), we calculate three sums of the edge weights.

1. The total sum of edge weights for all edges with features that are annotated to one or more of the annotations (*annotated*).
2. The *within* annotation edge weight sum, where both start and end nodes are annotated to the same annotation.
3. The *outer* annotation edge weight sum, where the start node is part of the annotated set, and the end node is annotated to one of the other annotations.

The partitioning ratio (or q-ratio) is calculated as follows:

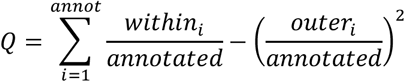

The partitioning ratio was originally designed as a method to determine the optimal clustering of networks, where each member of the network has only a single label [37–40]. In those cases, the partitioning ratio should range between 0 and 1 for non-partitioned and fully partitioned networks, respectively. The grouped Reactome pathways still have shared metabolite features, and therefore the partitioning ratios have a much wider range of values. However, we expect that **better** partitioning of the network will be reflected by a **more positive** partitioning ratios.

Partitioning ratios were compared across correlation methods using a paired t-test, where methods were paired by the dataset. P-values were adjusted using the Bonferroni correction.

### 2.14 Changes In Correlation Due to Changes in Dynamic Range and Imputation

We created a simulated dataset with 1000 features and 100 samples, starting with a sample from a random log-normal distribution with a mean-log value of 1, and standard deviation of 0.5. Uniform noise was added via random normal distribution with a standard deviation of 0.2 to create 100 samples from the base distribution, values were transformed to normal space and then *log*_10_ applied to have a representation of orders of magnitude and dynamic range. For any maximum level of censoring to be applied, a uniform distribution sample is generated on the range of 0 − *max* with 100 values. Data censoring was applied by taking the minimum observed value for a sample, and adding the censoring value from the uniform distribution. Any values in the sample below the censoring value are set to missing (NA).

Correlations were calculated between samples when no missing values are present (*reference*), and then again after censoring (*trimmed*). Two different correlation methods were used: ICI-Kt; and Pearson correlation. Imputation for Pearson correlation involved replacing all missing values with ½ the lowest observed value in the dataset after censoring. Differences between the reference and trimmed correlations were calculated, as well as the difference in the absolute value of differences of ICI-Kt and Pearson imputed.

### 2.15 Data Processing

All data processing and statistical analysis used R v 4.4.1 [22] and Bioconductor v 3.20 [41]. JSONized MW files were read in using jsonlite v 2.0.0 [42]. Metabolite feature id and sample id cleaning used janitor v 2.2.1 [43] and dplyr v 1.1.4 [44]. Plots were generated using either singly or in combination the packages: ggplot2 v 4.0.0 [45]; ggforce v 0.5.0 [46]; patchwork v 1.3.2 [47]. Analysis workflows were coordinated using targets v 1.11.4 [48].

## 3. Results

### 3.1 Datasets

We are aware of only one previous investigation of the causes of missingness in metabolomics datasets [37]. In Do et al. [37], the authors showed that there was a limit of detection (LOD) effect, with a dependence on the day the samples were run. Unfortunately, the KORA4 metabolomics dataset from Do et al. is not publicly available, so we could not attempt to redo their analysis of missing values with the same dataset.

Given the number of projects and analyses available in Metabolomics Workbench (MW), we sought to obtain a large number of individual datasets (MW analyses) from MW to evaluate both the phenomenon of left censorship and the information-content-informed Kendall-tau methodology.

In Figure 2, we provide a summary of the number of datasets remaining after filtering for various attributes (see Methods). From the starting 6501, we were able to retain 711 for this study. There were 1636 datasets that had no metabolite abundance data, or the metabolite data was not easily parseable or downloadable from external text files. A further 2162 datasets were excluded due to having < 100 metabolites. 755 were removed because they had < 5 samples in at least one group of subject sample factors (SSF) or < 2 SSF. 72 remaining datasets had a maximum intensity < 20, indicating either being log-transformed or a very small dynamic range. Finally, 769 datasets were removed due to either having no missing values, or the calculation of a correlation of rank with number of missing values across samples returned an NA value.

**Figure 2.**
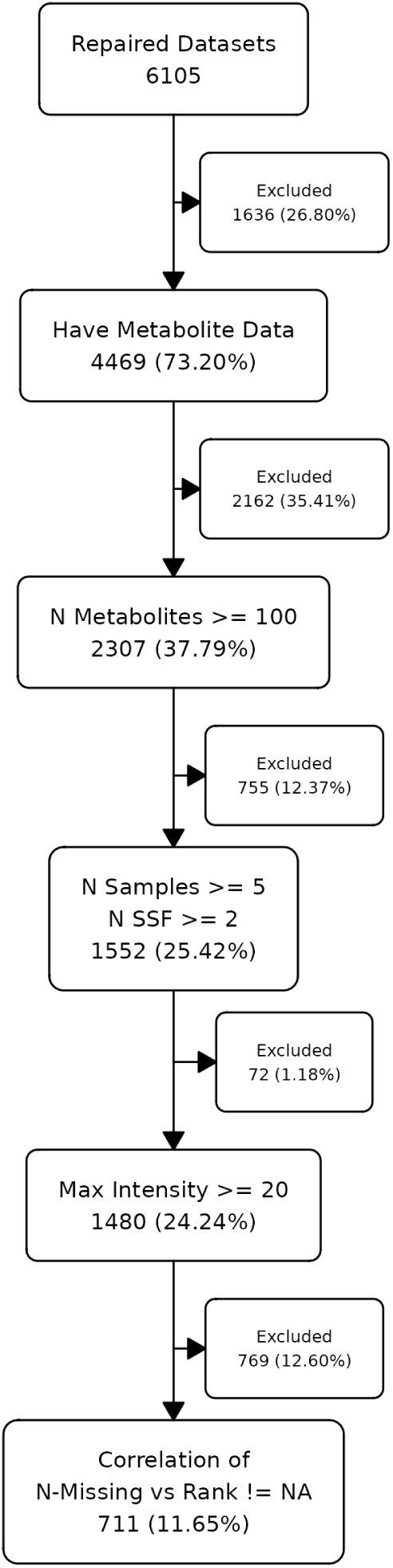
Number of datasets remaining after each level of filtering and / or checking.

### 3.2 Left Censoring As a Cause for Missingness

One would wonder just how many missing values are present in metabolomics datasets, and if their missingness is primarily due to left censorship or some other phenomenon. For the 711 datasets examined in this work, the percentage of missingness ranged from near zero (we required at least one missing value to keep a dataset for further analysis) to 25% for nuclear magnetic resonance (NMR), and the majority of mass-spectrometry (MS) datasets had missingness in the 0 - 25% range, with some datasets having missingness > 80% (Figure 3A). Using a binomial test to check if missing values are more likely to have non-missing values ranked at ≤ 0.5 of the values present, we see that the vast majority (681 of 711) have an adjusted-p-value ≤ 0.05, with over 160 having an extreme adjusted p-value (Figure 3B). We also checked if there is any relationship between the percentage of missing values and the binomial test p-values, but found none (Figure S1).

**Figure 3.**
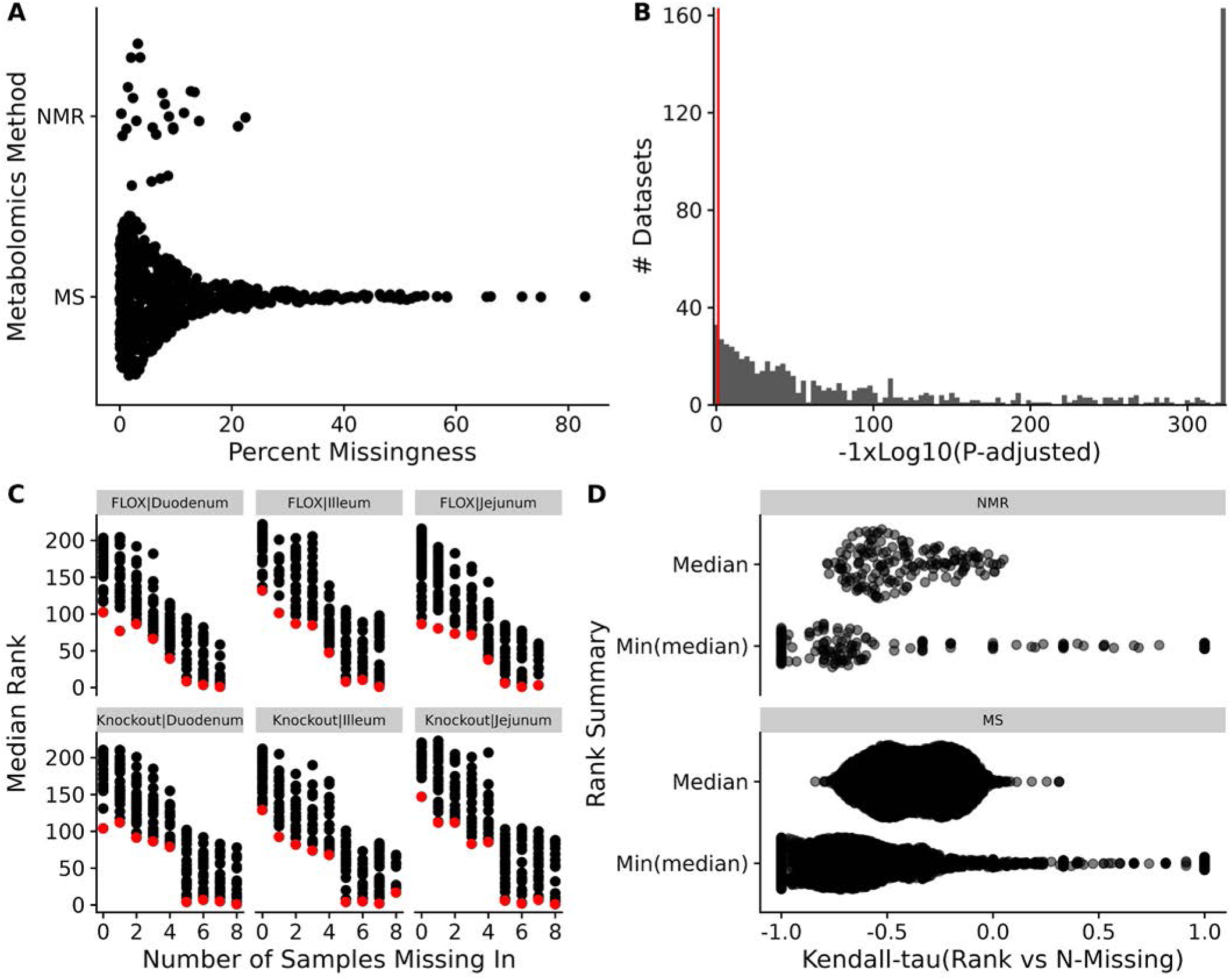
Missingness in datasets. **A**: Sina plot of the percent missing values in NMR and MS datasets. **B**: Histogram of log10 adjusted p-values from the binomial test testing if non-missing values are more likely to be below the median rank when the metabolite has missingness in the dataset. Red line indicates an adjusted p-value of 0.05. Adjusted p-values of 0 were replaced with the lowest observed non-zero adjusted p-value. **C**: Median (black) and minimum median (red) rank of metabolite abundances across factor groups of samples *vs* the number of samples the metabolite had a missing value for dataset AN001074. **D**: Sina plots of the Kendall-tau correlation of the median rank and minimum median rank with number of missing samples across all datasets.

For each set of subject sample factors (SSF) of a dataset, we calculated the median rank and number of measurement values missing across samples (i.e. N-Missing) for each metabolite. As shown in Figure 3C, there is a monotonically decreasing relationship between the median rank and the number of missing values for that metabolite. Moreover, as N-Missing decreases, there is clearly a minimum median rank below which the values do not cross as illustrated by the red points (Figure 3C). As shown in Figure 3D, median rank and N-missing are negatively correlated across the majority of experiments, although there are more positive correlations when using the minimum median rank. Given the results of the binomial test of missing ranks and this relationship of the minimum median value observed and N-Present, we believe that the **majority** of missing values in many metabolomics datasets are due to left censorship.

This makes the ICI-Kt methodology appropriate for use in many metabolomics datasets containing missing values through incorporation of the missing values below the LOD as useful information in the correlation calculation.

### 3.3 Comparison To Other Correlation Measures

We compared the ICI-Kt correlation to both Pearson and regular Kendall-tau-b correlations as calculated by the built-in R functions using simulated data sets with missing values (Figures S2-S4).

We created two samples with 1000 observations each drawn from a log-normal distribution, added further variance using a uniform distribution, and sorted in each case to create a pair of X and Y samples with a correlation of 1 and -1 for both Pearson and Kendall-tau correlation measures. The *true* correlation for each of the Kendall and Pearson correlations were calculated, and then missingness was introduced in the lower range of values, up to half of the values (see Simulated Data Sets).

In Figure 4 (and Figure S5), we can see that as missing values are added, only the ICIKt correlation values change in any significant way as illustrated by the wider range of ICI-Kt values on the y-axes versus the much narrower range of Pearson and Kendall tau correlation values on the x-axes. As the number of missing values increase, the ICI-Kt values drop or increase by up to 0.2. Similarly, Pearson correlation is also affected, but the degree of change in the correlation values are much less (notice the orders of magnitude differences in the x-axis scales compared to the y-axis), on the order of only 0.005 for both cases.

**Figure 4.**
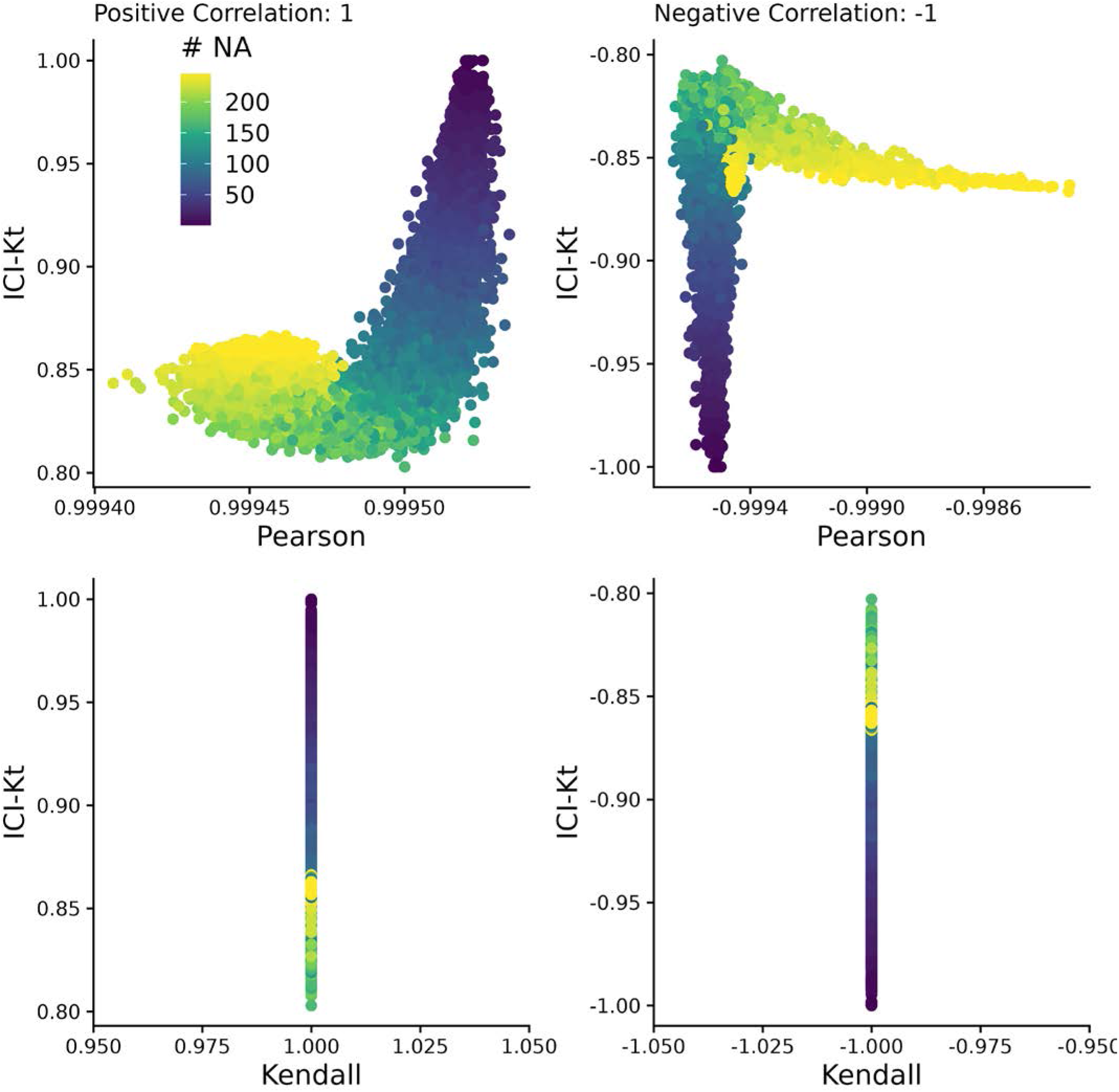
Comparing the correlation values obtained by Pearson, Kendall, and ICI-Kt correlation as an increasing number of missing values (0 - 500) in the bottom half of either sample for both positively (correlation = 1) and negatively (correlation = -1) correlated samples. Points are colored by how many points were set to missing on average between the two samples. A subset of 10,000 points was used for visualization.

Adding outlier points at the high end of the distribution to one of the samples causes very odd discrete patterns to appear in the negative Pearson correlations (see Figures S6, S7). Again, the scale of differences is much smaller in the Pearson correlations versus ICI-Kt. The negative Kendall correlations are unaffected by the outliers, in large part due to being a rank-based correlation. Likewise, the negative ICI-Kt correlations appear unaffected; however, the scale of changes seen in the negative Pearson correlations if present in the negative ICI-Kt correlations might simply be obscured by the changes due to missingness that are orders of magnitude larger.

These results demonstrate that the ICI-Kt correlation has quantitative sensitivity to missing values over the normal Kendall-tau correlation and linear Pearson correlation where points with missing values are ignored (pairwise complete).

### 3.4 Effect of Left Censoring VS Random Missing Data

Figure 5 demonstrates the effect of introducing left-censored versus random missingness in five different measures of correlation, including the ICI-Kt, the normal Kendalltau with *pairwise-complete-observations*, the normal Kendall-tau replacing missing with 0, Pearson with *pairwise-complete-observations*, and Pearson replacing missing with 0. The ICI-Kt correlation demonstrates a slight increase from the starting 0.90 correlation value with growing left-centered missingness caused by a slight reinforcement of the correlation, while with **growing random missingness**, the ICI-Kt correlation drops precipitously due to the large increase in discordant pairs caused by the random missing values. The normal Kendall tau correlation with pairwise complete has a small decrease in the correlation value with growing left-centered missingness caused by a loss of supporting pairs, while this correlation has a near constant average value with growing random missingness. The normal Kendall tau correlation replacing missing with 0 has identical behavior to the ICI-Kt correlation. In contrast to ICI-Kt, the Pearson correlation calculated using only pairwise complete entries is **practically constant** (i.e., range of 0.004 or less) over growing left-centered and random missingness. When replacing missing values with zero, Pearson correlation demonstrates a small decrease in the correlation value with growing left-centered missingness due to the zero values causing some deviation from linearity. Pearson correlation drops precipitously with growing random missingness with a magnitude similar to the ICI-Kt and normal Kendall tau replacing missing with 0. Overall, the ICI-Kt and the normal Kendall-tau replacing missing with zero have the desirable characteristics of maintaining the correlation with growing left-centered missing **while sharply dropping the correlation** with growing random missingness. In this special case where zero is lower than all of the values in the dataset, ICI-Kt and Kendall-tau replaced with zero result in identical correlation values, as shown in the bottom panels of Figure S8A and Figure S8C. In a naive treatment of the left-centered missing data, if the values below the cutoff are set to missing followed by log-transforming the values and subsequently setting missing values to 0, then the Kendall tau correlation replacing missing with 0 will show some very odd patterns at low intensity cutoffs due to the introduction of discordant pairs. Likewise, Pearson correlation replacing missing with 0 shows a parabolic effect with increasing missing values.

**Figure 5.**
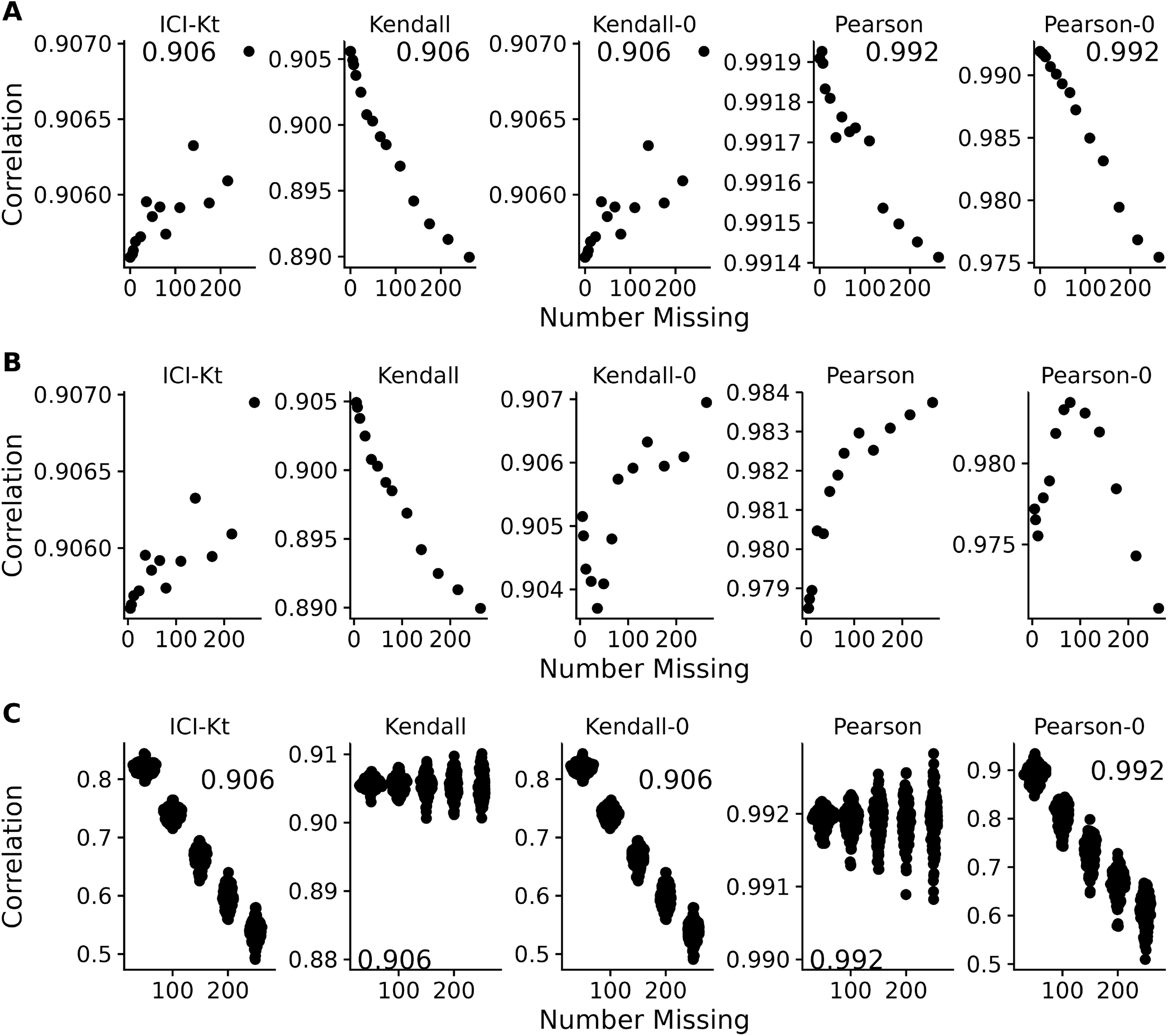
Effect of introducing missing values from a cutoff (**A** & **B**) or randomly (**C**) on different measures of correlation, including ICI-Kt, Kendall with pairwise complete, Kendall replacing missing with 0, Pearson with pairwise complete, and Pearson replacing missing with 0. **A**: Missing values introduced by setting an increasing cutoff. **B**: Missing values introduced by setting an increasing cutoff, and then log-transforming the values before calculating correlation. **C**: Missing values introduced at random. For the random case, each sample of random positions was repeated 100 times. Pay attention to the different y-axis ranges across graphs, with **A** and **B** graphs having much smaller y-axis ranges compared to **C**.

A common way missing data is handled in correlation calculations is to ignore them completely and use the pairwise complete cases to calculate the Pearson correlation coefficient. As shown in Figure 5C, this results in a complete misestimation of the changed correlative structure introduced by random entries. ICI-Kt, in contrast, incorporates the missingness in a sensical way, and the resulting correlation values fall as random entries are introduced.

### 3.5 Differences in Dynamic Range and Correlation

Another way that missing values appear is due to changes in dynamic range between samples, as some samples have features with higher values, and the fixed dynamic range of the instrumentation results in features with lower values to be “missing” in those samples. We created a set of 100 simulated samples with uniform noise on the log-scale, with relatively constant dynamic ranges, and introduced changes to the overall dynamic range using a random censor at varying levels (see Implementation). Possible different levels of censoring based on dynamic range were checked by first determining how many missing values would be introduced in each sample as the dynamic range was increased in increments of 0.1 (see Figure S9). Based on the number of values being censored, limits of 0.5, 1, and 1.5 were selected, representing low, medium and high variability of the dynamic range.

For each level of possible missingness introduced by changes to the dynamic range, correlation across all samples were calculated using all values (reference), as well as after missingness was added (trimmed), and using Pearson correlation with global imputation (Pearson Imputed), or ICI-Kt. Figure 6 demonstrates that it is only as the number of missing values in one of the samples approaches 50% or more (500 of 1000 features) does the Pearson correlation with global imputation give correlation values closer to the known correlation with no missing values in any appreciable amount (points below the red lines in the top panels, and to the right of the red line in the histograms in the bottom panels). Points above the lines with slopes of -1 and 1 indicate that the difference of reference - trimmed is smaller in the ICI-Kt correlations, and below the lines indicate the difference is larger in the ICI-Kt correlations. This is further emphasized by the majority of the values are to the left of the line at 0 in the difference histograms.

**Figure 6.**
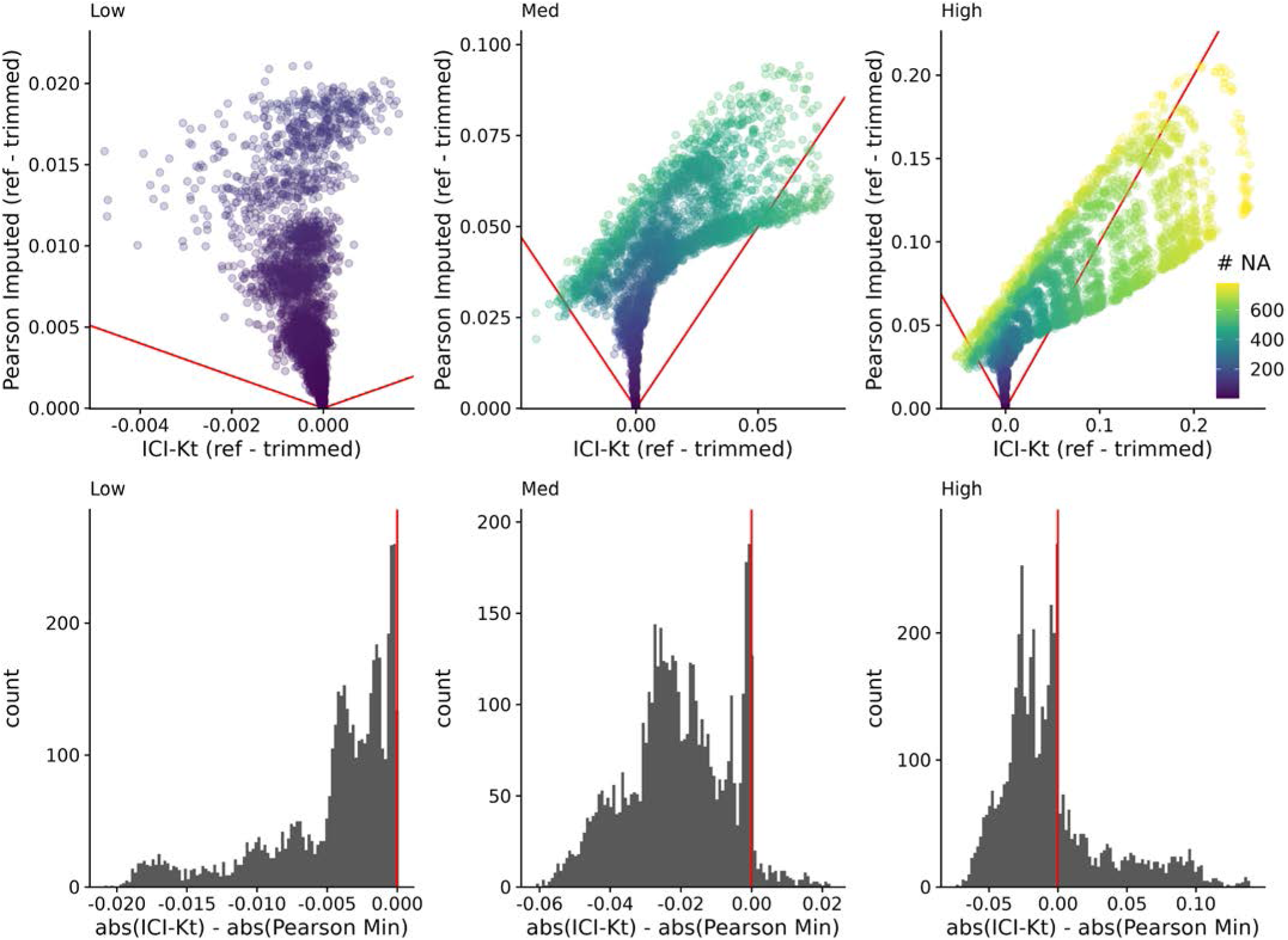
Top: Difference of reference - trimmed ICI-Kt correlation *vs* Pearson imputed using ½ the minimum value in the dataset. Low, med, and high indicate the level of variability in dynamic range, using 0.5, 1, and 1.5, respectively. Red lines indicate slope of -1 and 1. Color indicates the maximum number of missing values between the two samples being correlated. Bottom: Differences in the absolute value of reference - trimmed differences between ICI-Kt and Pearson imputed correlations.

### 3.6 Utility for Metabolomics Data Sets

Having established that many metabolomics data sets with missing values are present due to left-censorship (see Figure 3), we analyzed how the ICI-Kt methodology compares to other methods for outlier removal and for the generation of feature-feature networks.

In outlier removal, the evaluation used differential analysis of metabolites across conditions. For each correlation method, outlier samples within each SSF were determined and removed, and an F-test conducted using the limma package across SSF. The fraction of metabolites that were differential after outlier removal was determined, and a t-test used to evaluate pairwise comparisons of methods to see if any differences were significant.

Table 1 shows the fractions of significant metabolites after removing outlier samples using each method. Table 2 shows the statistical results of the pairwise comparisons of each method based on the significant fractions. Both tables show that while there is a significant change in the fraction of significant metabolites after removal of outlier samples, the actual average differences are very small.

**Table 1.** Summary statistics of the fraction of significantly different features after removing outliers detected using each correlation method.

| Method | Mean | SD | Median | MAD |
| --- | --- | --- | --- | --- |
| icikt | 0.457 | 0.338 | 0.454 | 0.482 |
| icikt_complete | 0.457 | 0.336 | 0.450 | 0.480 |
| pearson_log1p | 0.455 | 0.337 | 0.450 | 0.487 |
| kt_base | 0.455 | 0.337 | 0.460 | 0.478 |
| pearson_log | 0.453 | 0.337 | 0.450 | 0.475 |
| pearson_base | 0.450 | 0.338 | 0.442 | 0.483 |
| pearson_base_nozero | 0.448 | 0.338 | 0.441 | 0.480 |
| original | 0.443 | 0.338 | 0.436 | 0.484 |

**Table 2.** Paired t-test statistical results comparing the fraction of significant metabolite features after removing outlier samples using different methods. Adjusted p-values were calculated using the Bonferroni method.

| Comparison | Difference | P-Value | P-Adjusted |
| --- | --- | --- | --- |
| icikt v original | 0.0137 | 7.9e-13 | 3.5e-11 |
| icikt_complete v original | 0.013 | 1.5e-11 | 6.9e-10 |
| kt_base v original | 0.0104 | 1.1e-09 | 4.9e-08 |
| pearson_log1p v original | 0.011 | 2.7e-09 | 1.2e-07 |
| pearson_log v original | 0.00964 | 4.1e-09 | 1.8e-07 |
| icikt v pearson_base | 0.00819 | 3.8e-08 | 1.7e-06 |
| icikt v pearson_base_nozero | 0.00949 | 5.5e-08 | 2.5e-06 |
| icikt_complete v pearson_base_nozero | 0.00888 | 1.5e-06 | 6.8e-05 |
| icikt_complete v pearson_base | 0.00758 | 3.6e-06 | 1.6e-04 |
| pearson_base_nozero v kt_base | -0.00625 | 2.7e-05 | 1.2e-03 |
| pearson_base_nozero v pearson_log1p | -0.00687 | 1.0e-04 | 4.7e-03 |
| pearson_base_nozero v pearson_log | -0.00547 | 2.0e-04 | 9.0e-03 |
| pearson_base v kt_base | -0.00495 | 2.3e-04 | 1.0e-02 |
| pearson_base v original | 0.00547 | 2.6e-04 | 1.2e-02 |
| pearson_base v pearson_log1p | -0.00558 | 2.8e-04 | 1.3e-02 |

In feature-feature network generation, we evaluated the differences in partitioning ratios of metabolite features across aggregated Reactome pathways, after creating weighted feature-feature networks using the various correlation methods. Paired t-tests compared the methods, and are reported in Table 3 (and Table S2), and graphed in Figure S10. Both the ICI-Kt complete and base variant show much larger positive differences in partitioning ratio compared to all other methods, including the base Kendall-tau. This implies that the gains in partitioning ratio are not only due to using a rank-based correlation.

**Table 3.**
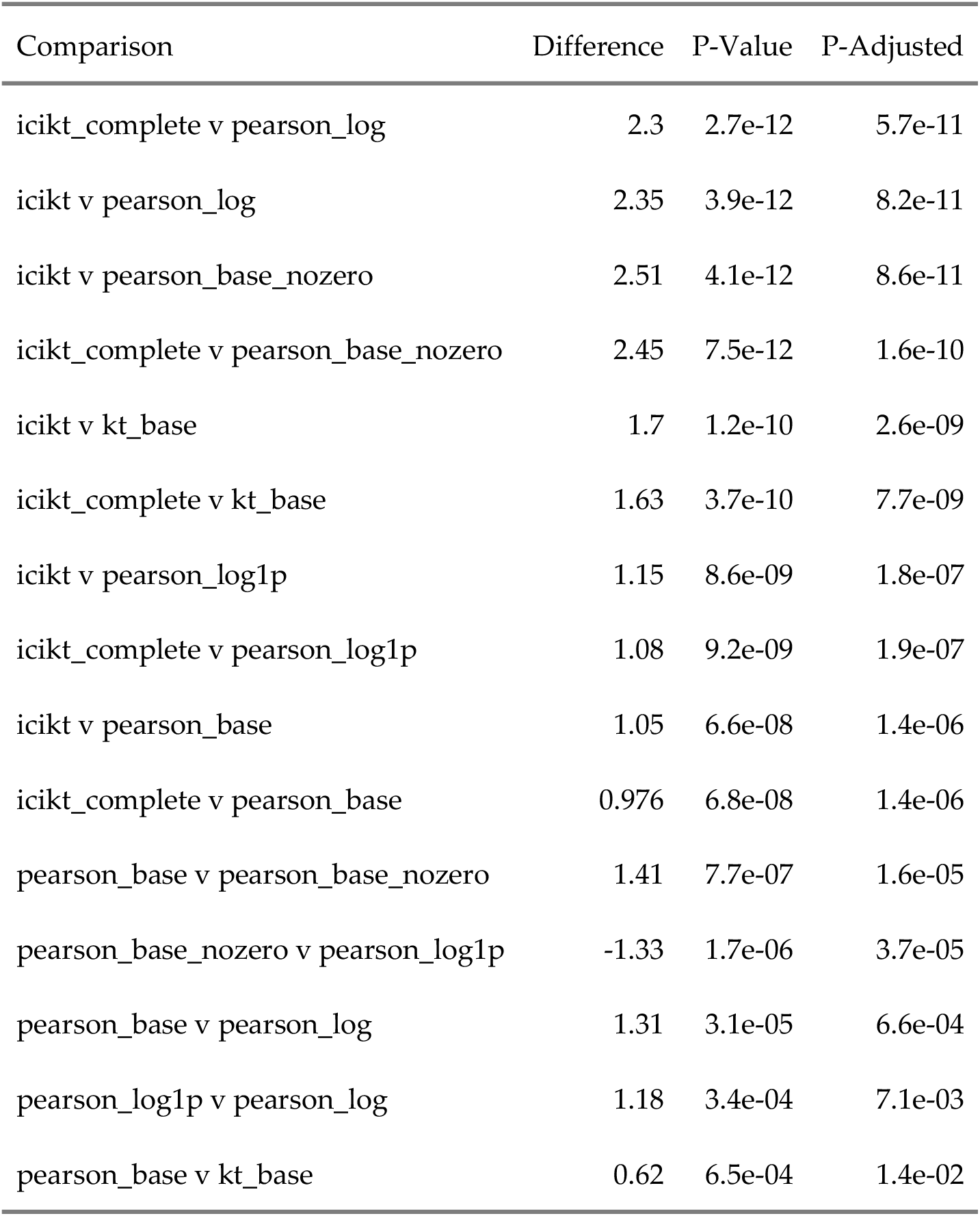
Paired t-test statistical results comparing the partitioning ratio of networks generated by the various correlation methods. Adjusted p-values were calculated using the Bonferroni method.

| Comparison | Difference | P-Value | P-Adjusted |
| --- | --- | --- | --- |
| icikt_complete v pearson_log | 2.3 | 2.7e-12 | 5.7e-11 |
| icikt v pearson_log | 2.35 | 3.9e-12 | 8.2e-11 |
| icikt v pearson_base_nozero | 2.51 | 4.1e-12 | 8.6e-11 |
| icikt_complete v pearson_base_nozero | 2.45 | 7.5e-12 | 1.6e-10 |
| icikt v kt_base | 1.7 | 1.2e-10 | 2.6e-09 |
| icikt_complete v kt_base | 1.63 | 3.7e-10 | 7.7e-09 |
| icikt v pearson_log1p | 1.15 | 8.6e-09 | 1.8e-07 |
| icikt_complete v pearson_log1p | 1.08 | 9.2e-09 | 1.9e-07 |
| icikt v pearson_base | 1.05 | 6.6e-08 | 1.4e-06 |
| icikt_complete v pearson_base | 0.976 | 6.8e-08 | 1.4e-06 |
| pearson_base v pearson_base_nozero | 1.41 | 7.7e-07 | 1.6e-05 |
| pearson_base_nozero v pearson_log1p | -1.33 | 1.7e-06 | 3.7e-05 |
| pearson_base v pearson_log | 1.31 | 3.1e-05 | 6.6e-04 |
| pearson_log1p v pearson_log | 1.18 | 3.4e-04 | 7.1e-03 |
| pearson_base v kt_base | 0.62 | 6.5e-04 | 1.4e-02 |

## 4 Discussion and Conclusion

Left-censored distributions in analytical measurements of biological samples are common in biological and biomedical research, because of detection limits of the analytical instrumentation, which produces missing measurement values for all the analytes below these detection limits. As far as we are aware, previous efforts in this area are concerned with either 1: attempting to come up with better imputation methods prior to calculating correlation values; or 2: finding ways to still estimate the correlation in the face of missing values, generally by finding maximum likelihood estimates. In the case of (1), there are many imputation methods, and new methods are still being developed, although they tend to be new combinations of old methods to handle the various types of missing values. For (2), the maximum likelihood methods generally apply to Pearson and similar types of correlation, as they benefit from the use of regression in the presence of missing data. Alvo and Cabilio’s work from 1995 [49] is one of the only works we are aware of that attempts to create a general framework for rank-based statistics in the presence of missing data. But, in our understanding their framework applies to data that is missing at random versus non-random missing-values, as is the case for analytes that are below the detection limit. Additionally, there does not appear to be a software implementation of Alvo and Cabilio’s method available.

Although the actual implementation of the base ICI-Kt correlation metric involves a global imputation of missing values, our equations demonstrate a left-censorship interpretation of missing values as useful information within the calculated correlation. Furthermore, the addition of “local”, “global” and *τ*_*max*_ normalizations of the ICI-Kt correlation in combination with completeness provide additional interpretations of information content. Finally, the availability of the binomial left-censorship test ensures the application of the ICI-Kt methodology when it is appropriate.

Global imputation methods rely on the assumption that samples have similar dynamic range of detection and thus an imputed value should be comparable between samples. However, the dynamic range of detection is often variable across samples. For complex analytical techniques often used in omics experiments, the variability in the dynamic range of detection can be quite high. Under these circumstances, the ICI-Kt method provides more robust results as compared to Pearson correlation with global imputation. This holds true for low, medium, and high variability in dynamic range across samples.

In the case of using sample-sample correlation to detect outliers, imputation does not solve any of the issues related to discovering outliers, as it should be applied **after** outlier samples are removed, otherwise the imputed values may not be useful. When used to create feature-feature networks based on partial correlations derived from the feature-feature correlations, ICI-Kt based methods showed the best partitioning of features based on predicted pathway annotations. As far as we know, information-content-informed Kendall-tau (ICI-Kt) is the first correlation method that explicitly attempts to utilize non-random missing values that occur due to being below the detection limit. ICI-Kt **explicitly** treats left-censored missing values as correlative information while preserving the full deleterious effects of missing at random values on the correlation metric. Moreover, ICI-Kt can be combined with measurement completeness to create a composite metric that is quite sensitive to overall data quality on a sample-specific level. Also, this ICI-Kt × completeness metric may have applications in cluster detection of single-cell omics data sets.

The implementations of the ICI-Kt method in the available R and Python packages provide a rich set of options for customizing the correlation calculation for a variety of use-cases and interpretations. These packages handle missing values in log-transformed data in a safe manner and have O(nlogn) performance, making them computationally practical for real-world omics data sets. Also, these packages provide multiprocessing implementations that take full advantage of modern multi-core central processing units.

As demonstrated with the datasets analyzed here, the “best” correlation-related metric will likely depend on the specific dataset and the specific data analysis step. Many factors affect this, especially correlation linearity and the modality of measurement value distributions. We would humbly suggest that for most metabolomics datasets, the application of several correlation-related metrics simultaneously would be the best approach for outlier detection in quality control and quality assessment steps. Where one metric lacks outlier detection sensitivity, another metric will prove sensitive. Therefore, ICI-Kt and associated composite metrics should be considered as useful additions to the omics data analysis toolkit.

## Author Contributions

RMF wrote the code in the ICIKendallTau R package, tested the ICI-Kt correlation code, and wrote all of the analysis code for this manuscript. PSB wrote the icikt Python package. HNBM conceived of the ICI-Kt correlation metric, provided input into code structures, and supervised the analyses and interpretation of results. All authors contributed to the writing of the manuscript.

## Availability Of Data And Materials

GitHub repository for the R package: https://github.com/moseleyBioinformaticsLab/ICIKendallTau. GitHub repository for the Python package: https://github.com/moseleyBioinformaticsLab/icikt. Python Package Index: https://pypi.org/project/icikt/. Code and data used in the results for this manuscript are available on Zenodo: https://zenodo.org/records/18625643.

## Acknowledgements

We are heavily indebted to the University of Kentucky Center for Computational Sciences (CCS) provided Kentucky Research Informatics Cloud (KyRIC), an NSF supported computational resource (NSF ACI1626364) for access to large virtual machines that allowed for methods development. We are deeply indebted to all those research groups who deposited data to The Metabolomics Workbench, https://www.metabolomicsworkbench.org/, and the work The Metabolomics Workbench does in archiving and making datasets available. We thank Travis Thompson for downloading and repairing The Metabolomics Workbench datasets using a prototype of the mwtab v 2.0 package and related software.

## Abbreviations

The following abbreviations are used in this manuscript:

ICI-Kt: Information-content-informed Kendall-tau
EDA: exploratory data analysis
PCA: principal component analysis
MW: The Metabolomics Workbench
SSF: subject sample factors
LOD: limit of detection
NMR: nuclear magnetic resonance
MS: mass-spectrometry

## Supplemental Materials

### Left Censoring as a Cause for Missingness

**Figure S1.**
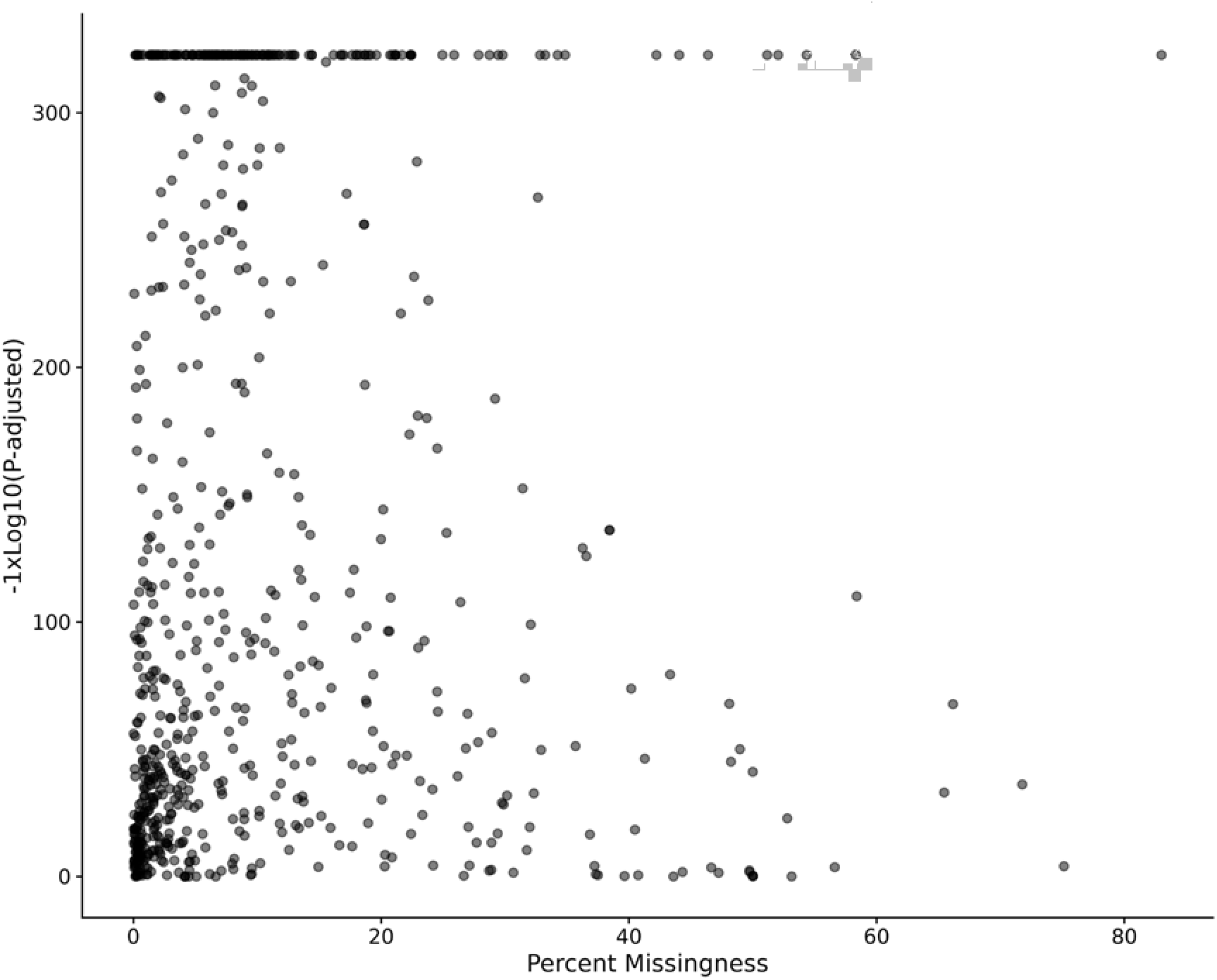
Percent missing values in each dataset compared to the p-values from the left censored binomial test.

### Simulated Data

**Figure S2.**
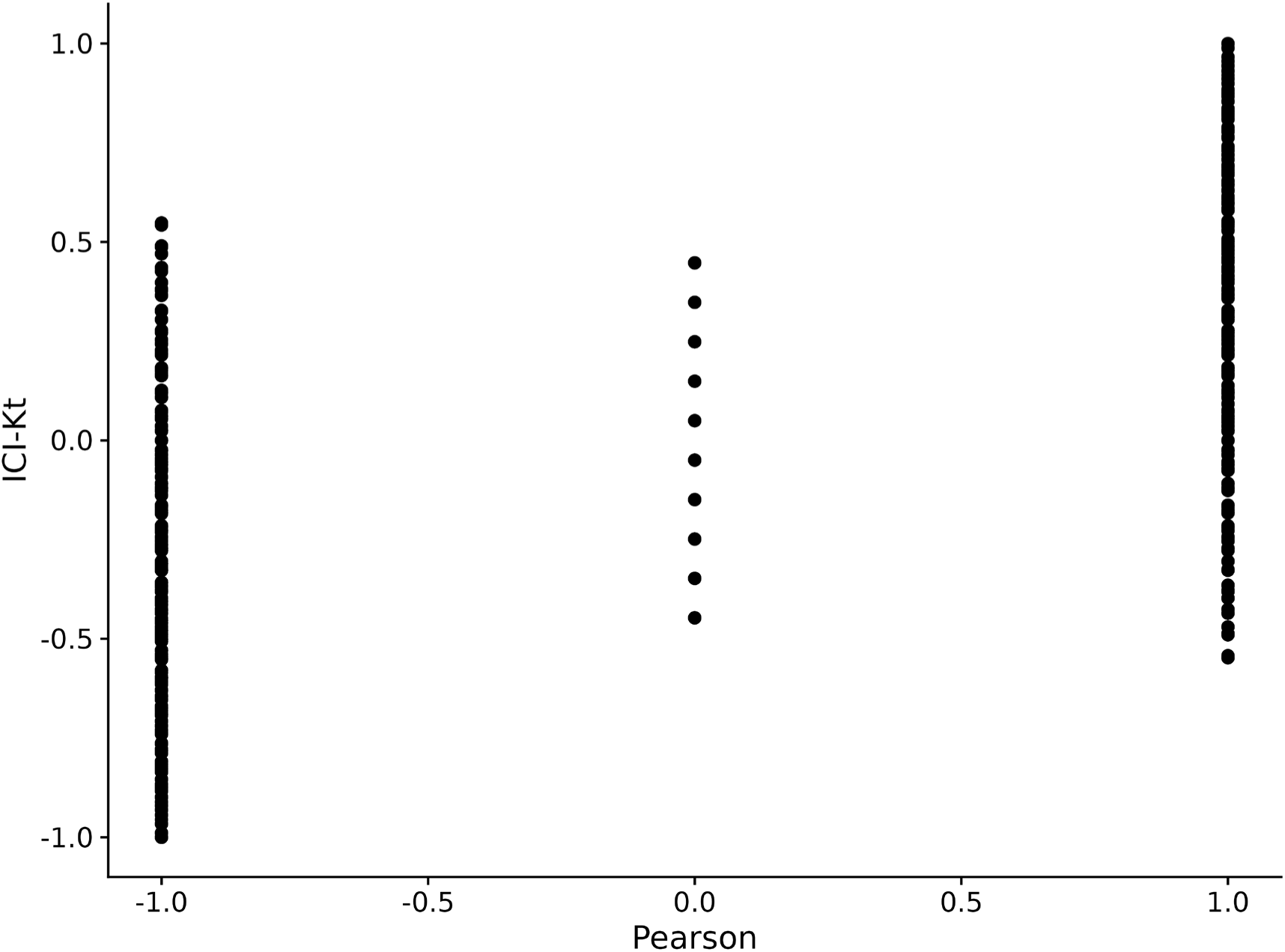
Comparison of ICI-Kt and Pearson correlations for perfectly positive and negatively correlated samples, systematically replacing values with NA. NA values from Pearson were replaced with zero for this comparison.

### Simple Data Set

We can also examine the full set of positive and negative correlations generated as we vary the number of missing entries between two positively correlated samples and two negatively correlated samples. These distributions are shown in Figure S4. We can see that the distributions from both ICI-Kt and Kendall-tau are the same, which is expected given we replaced missing values (NA) with zero *within* the ICI-Kt code, and replaced missing values (NA) with zero prior to calculating Kendall-tau correlations.

**Figure S3.**
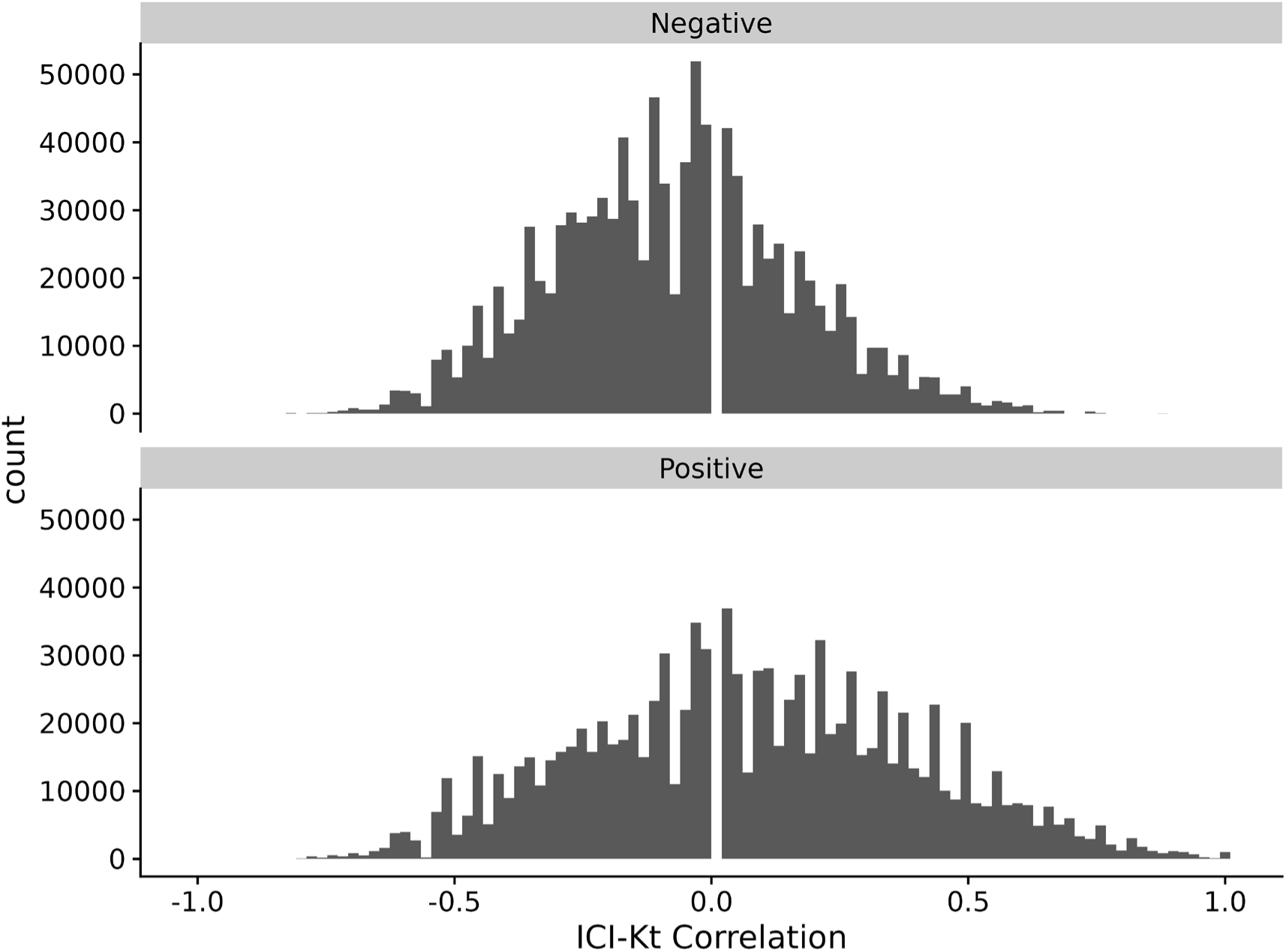
ICI-Kendall-tau correlation as missing values are varied between two samples.

**Figure S4.**
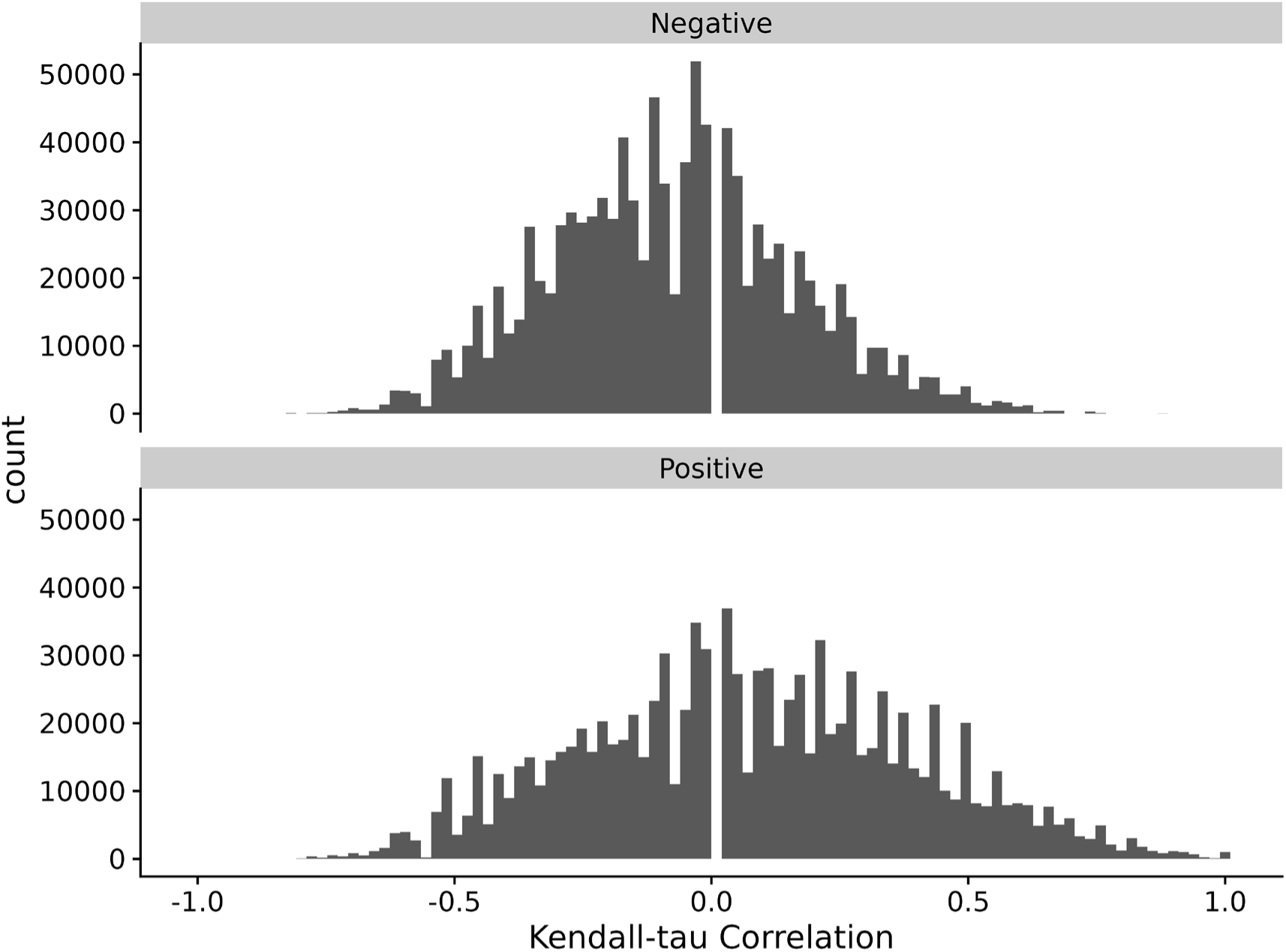
Kendall-tau correlation as missing values are varied between two samples and replaced with 0 before calculating Kendall-tau.

### Comparison to Other Correlation Measures

**Figure S5.**
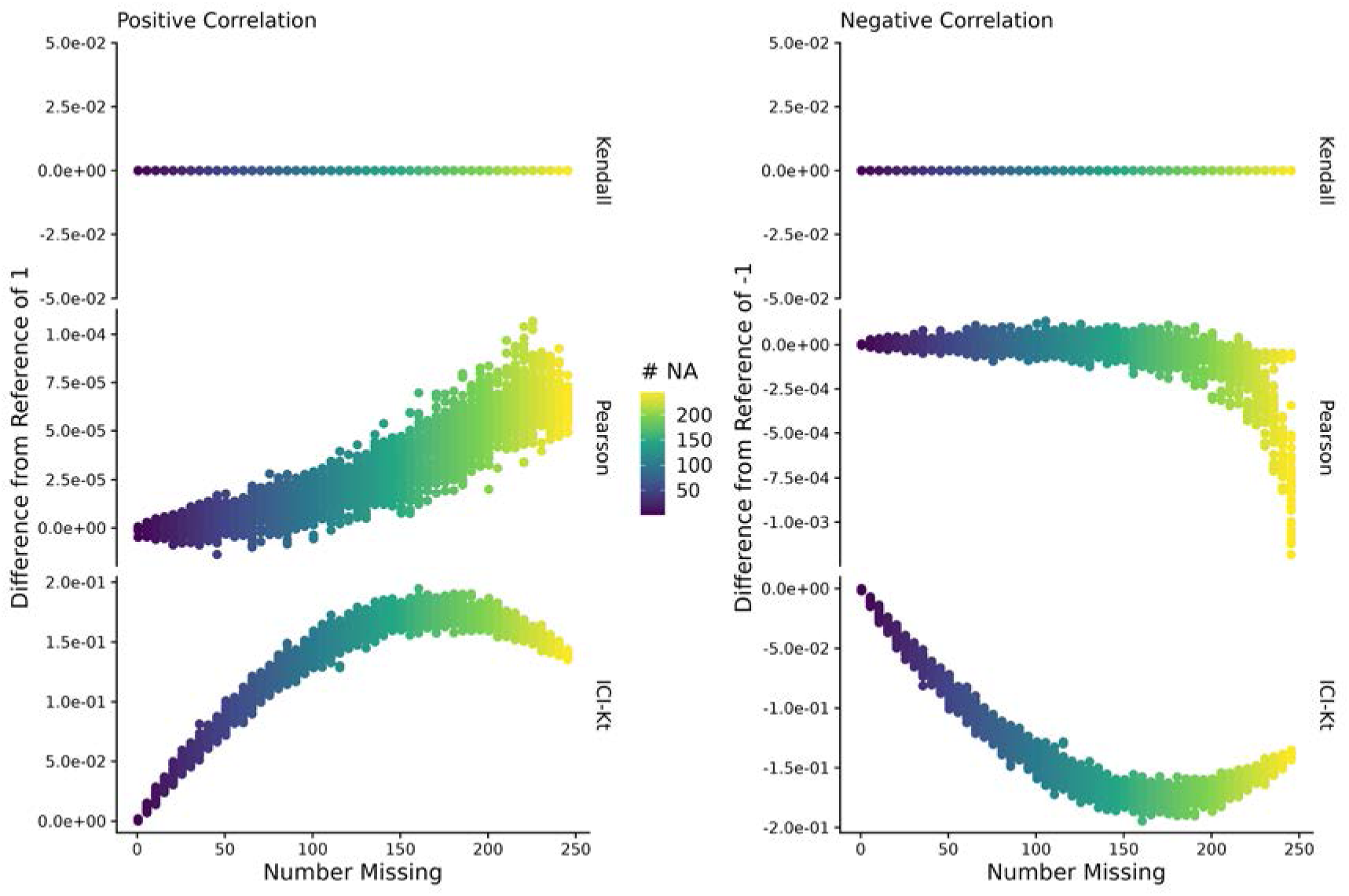
Difference of estimated correlation with missingness introduced compared to a reference correlation of 1 for the positive or -1 for the negative case, as a function of the average number of missing entries in X and Y sample (# NA). Points are colored by how many points are missing on average between the two samples X and Y.

**Figure S6.**
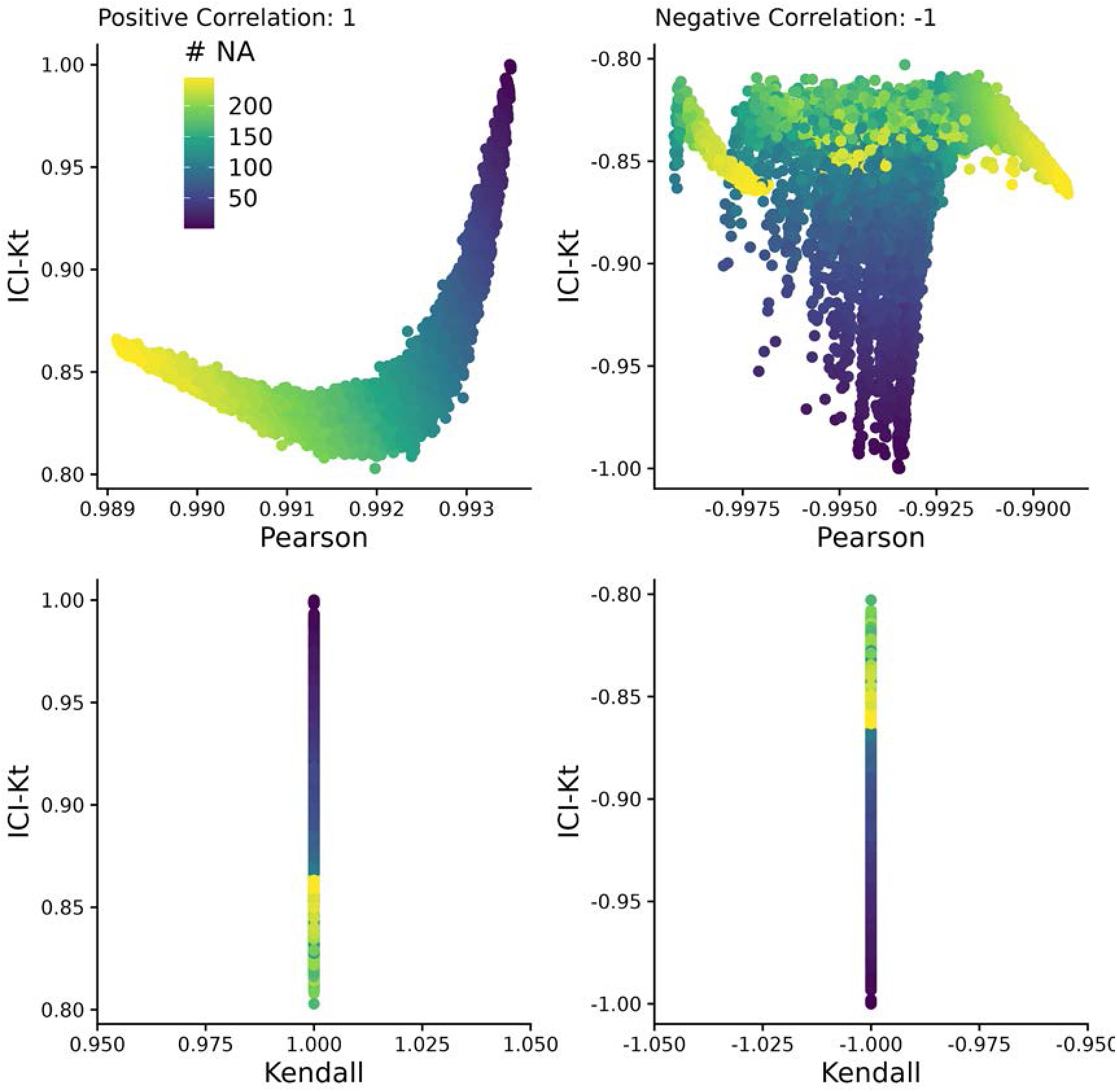
More variance was introduced at one end of the log-normal distribution in one sample to create a small percentage of outlier points compared to the other samples compared. This figure is comparing the correlation values obtained by Pearson, Kendall, and ICI-Kt correlation as an increasing number of missing values (0 - 500) in the bottom half of either sample for both positively (correlation = 1) and negatively (correlation = -1) correlated samples. Points are colored by how many points were set to missing on average between the two samples. A subset of 10,000 points was used for visualization.

**Figure S7.**
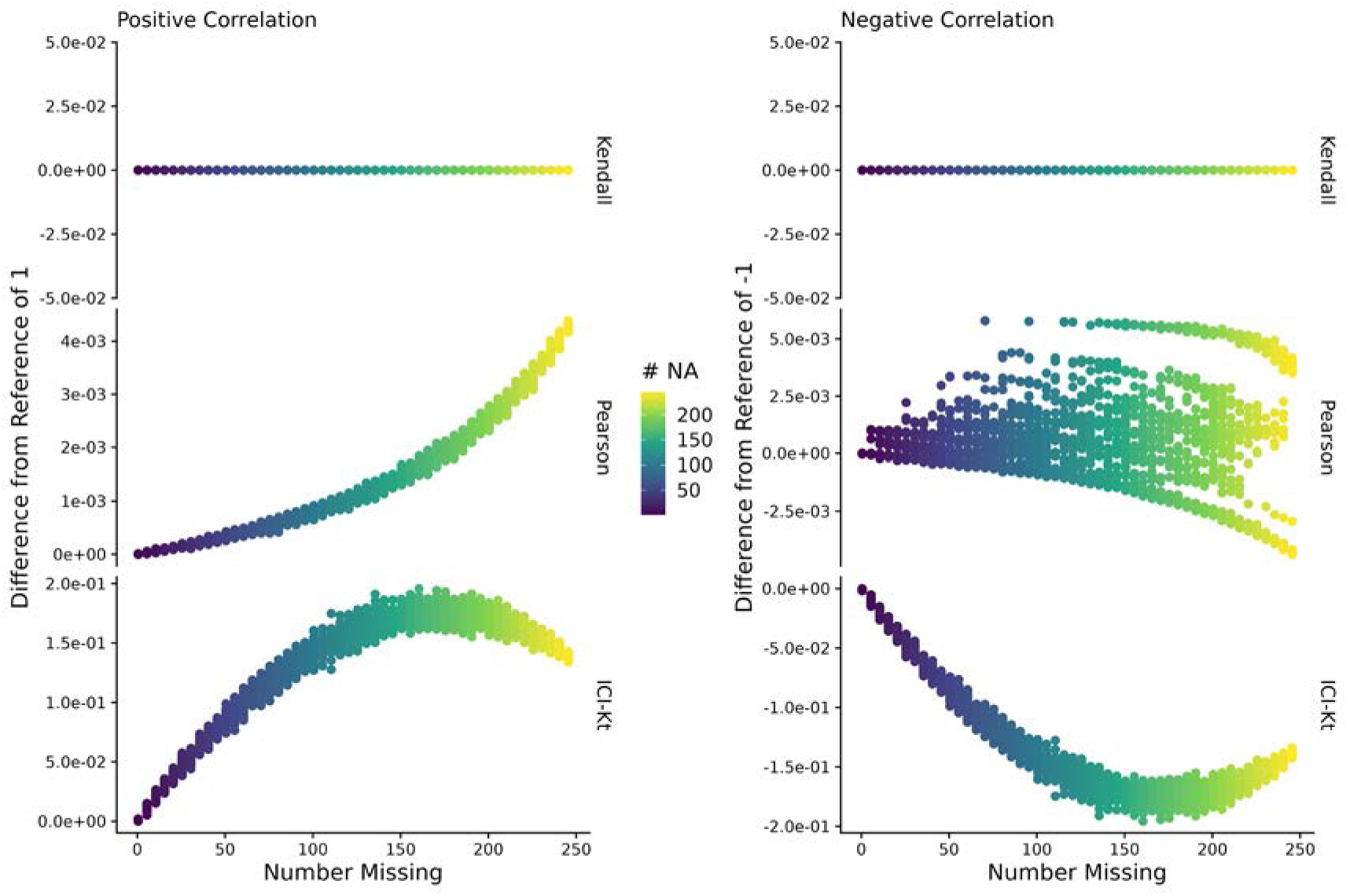
More variance was introduced at one end of the log-normal distribution in one sample to create a small percentage of outlier points compared to the other samples compared. This figure is showing the difference of estimated correlation with missingness introduced compared to a reference correlation of 1 for the positive or -1 for the negative case, as a function of the average number of missing entries in X and Y sample (# NA). Points are colored by how many points are missing on average between the two samples X and Y.

### Semi-Realistic Data Set

**Figure S8.**
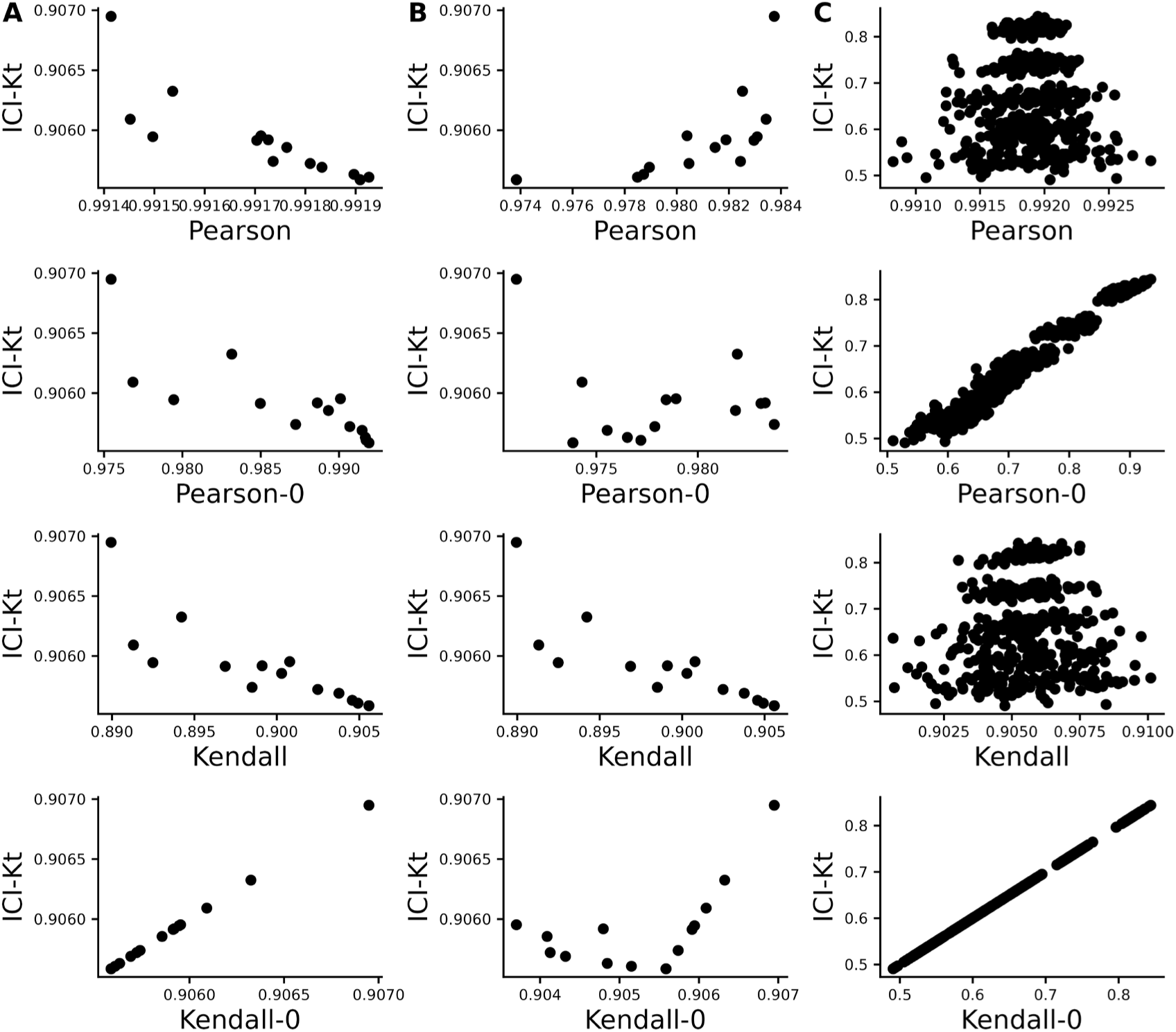
Effect of introducing missing values from a cutoff (A & B) or randomly (C) on different measures of correlation, including ICI-Kt, Kendall with pairwise complete, Kendall replacing missing with 0, Pearson with pairwise complete, and Pearson replacing missing with 0. A) Missing values introduced by setting an increasing cutoff. B) Missing values introduced by setting an increasing cutoff, and then log-transforming the values before calculating correlation. C) Missing values introduced at random. For the random case, each sample of random positions was repeated 100 times.

### Changes In Correlation Due to Changes in Dynamic Range and Imputation

**Figure S9.**
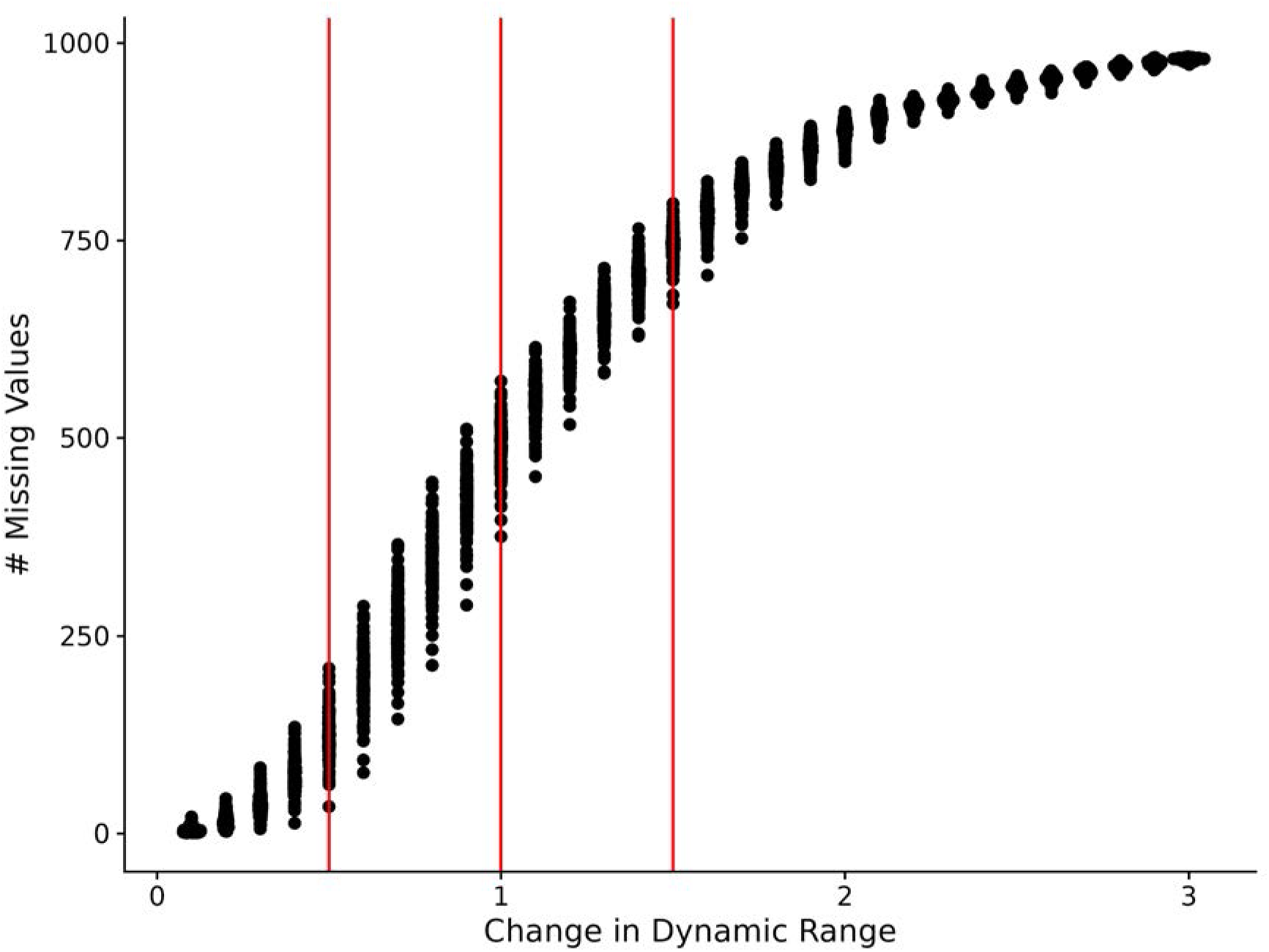
Number of missing values in each of the 100 samples as a function of changing the dynamic range of the values in the sample by increasing the lower limit of detection.

### Outlier Detection

**Table S1.**
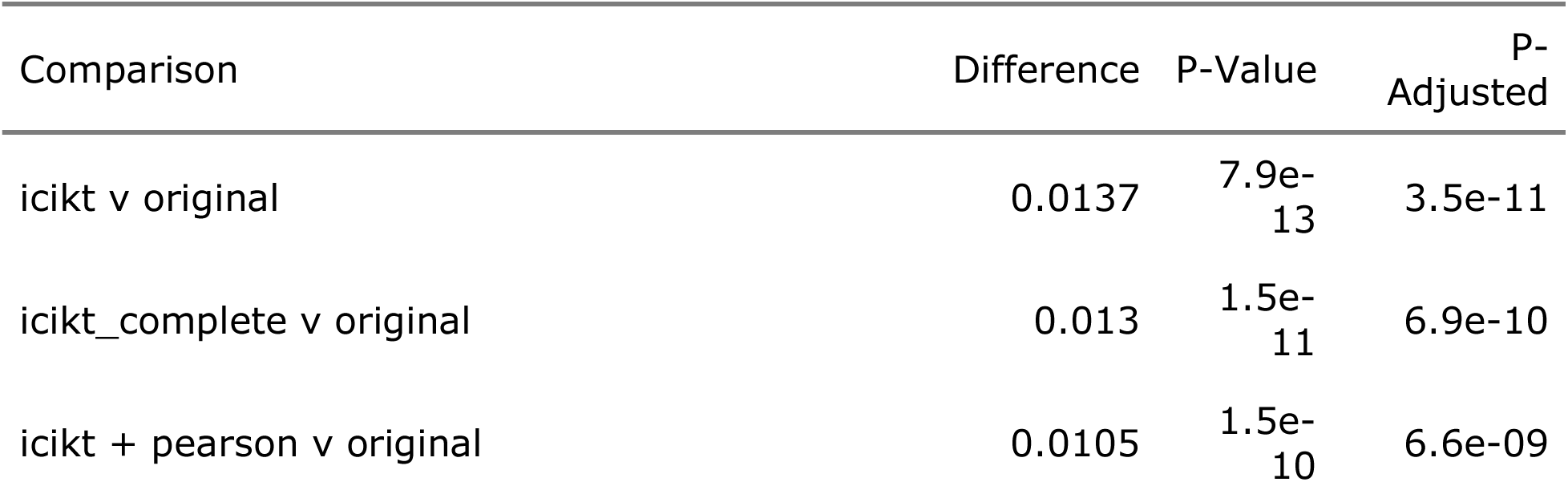
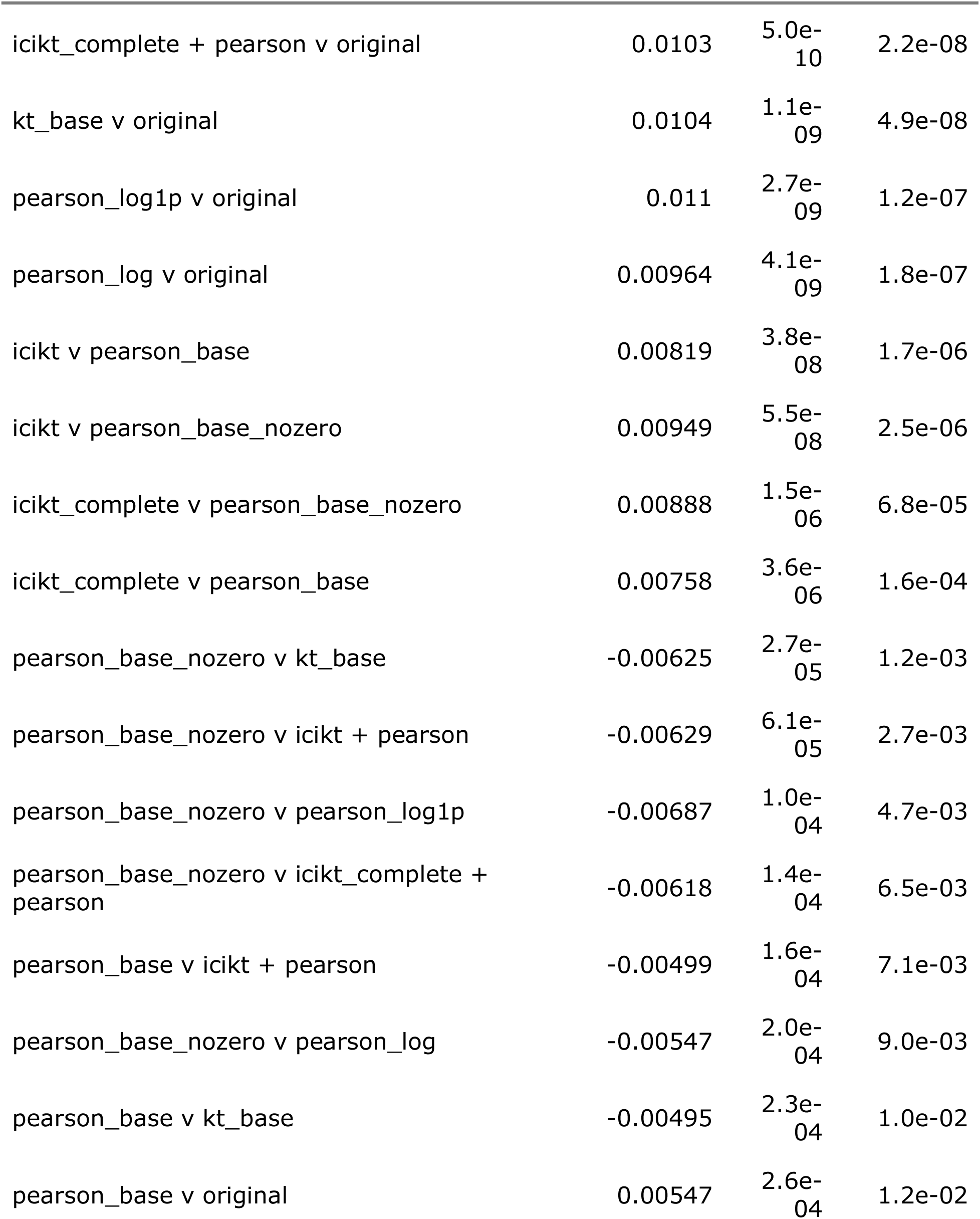
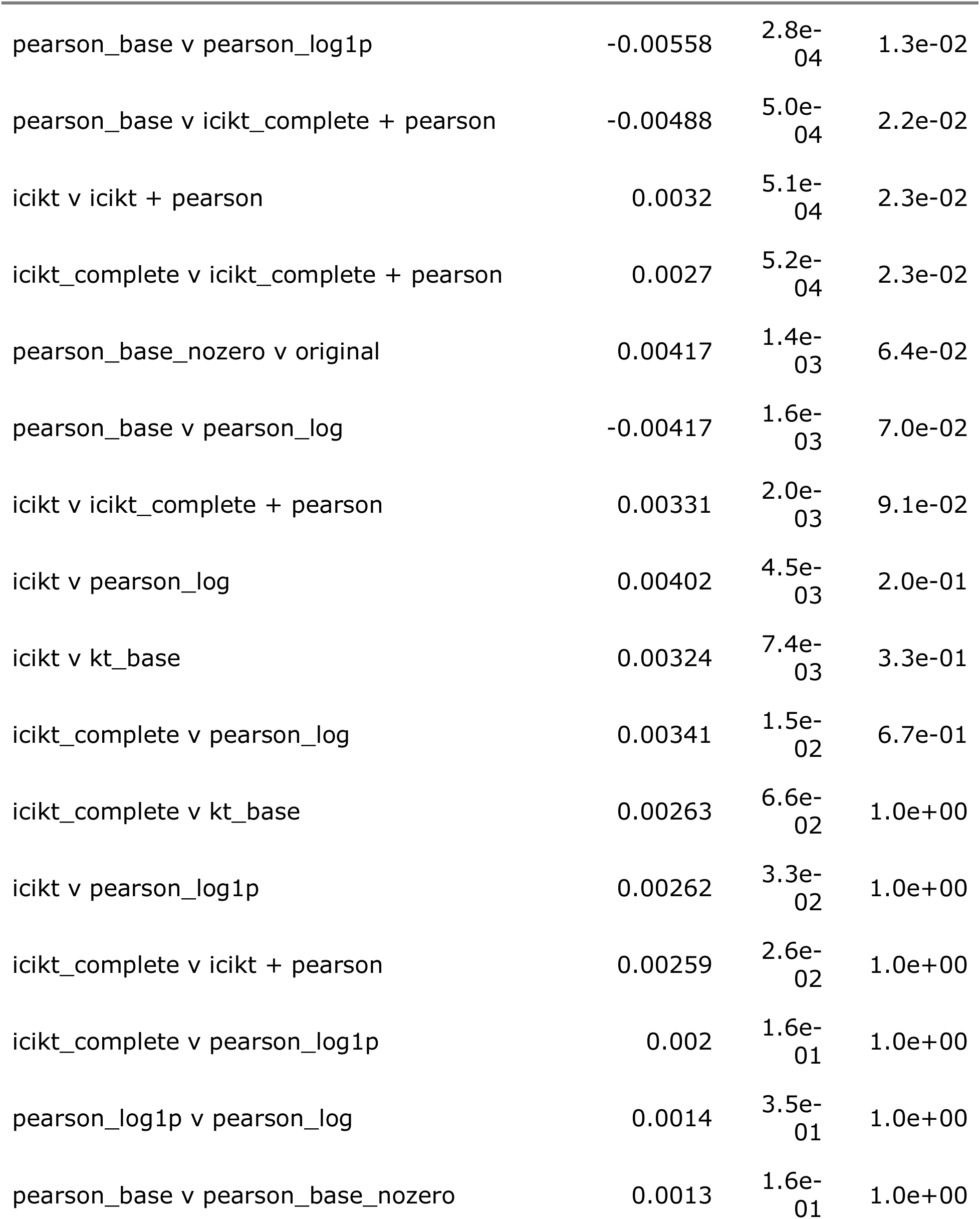
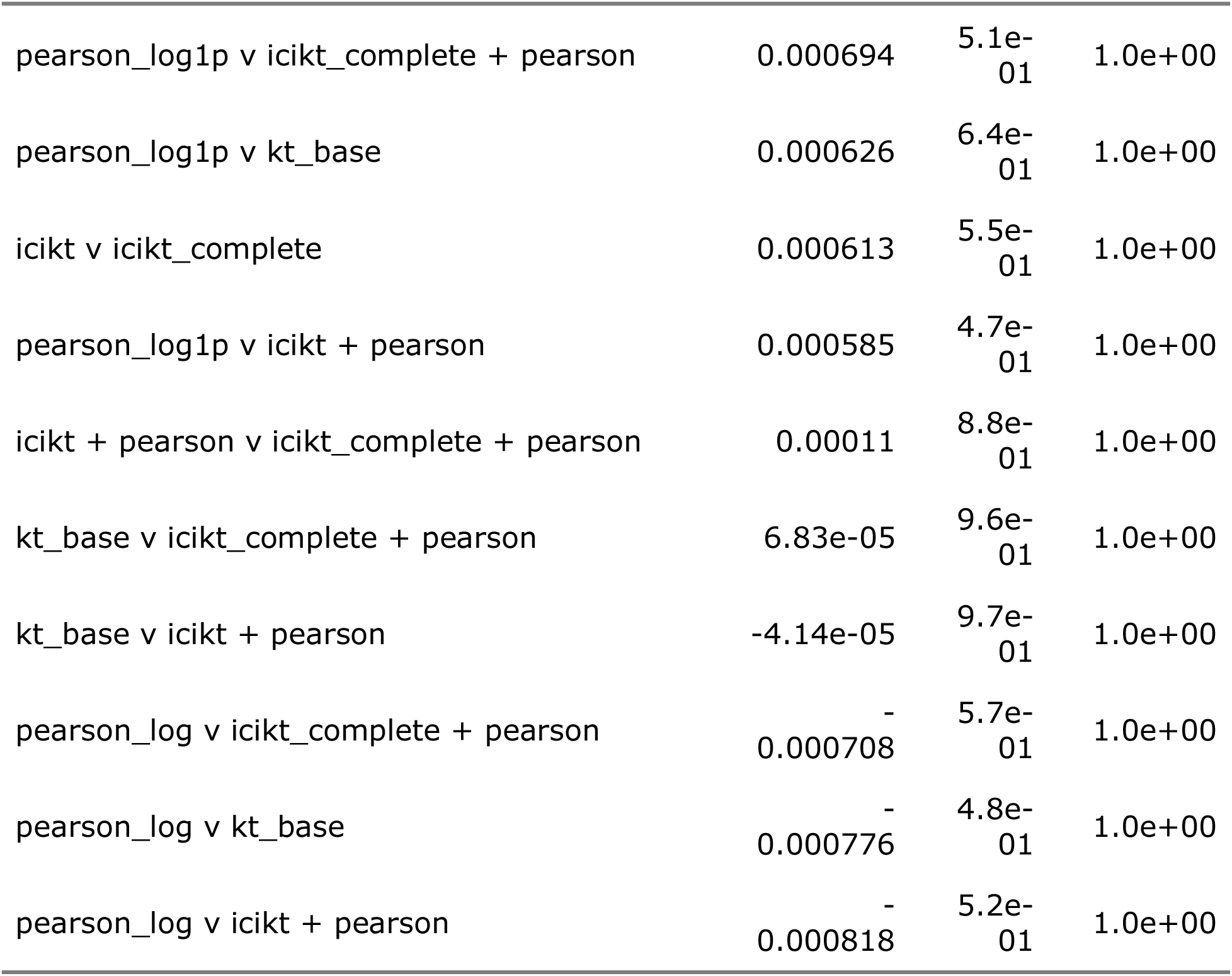
The statistical results of the pairwise comparisons of each method based on the significant fractions after removing outliers. P-values were adjusted using the Bonferroni method.

| Comparison | Difference | P-Value | P-Adjusted |
| --- | --- | --- | --- |
| icikt v original | 0.0137 | $7.9e-13$ | $3.5e-11$ |
| icikt_complete v original | 0.013 | $1.5e-11$ | $6.9e-10$ |
| icikt + pearson v original | 0.0105 | $1.5e-10$ | $6.6e-09$ |
| icikt_complete + pearson v original | 0.0103 | 5.0e-10 | 2.2e-08 |
| kt_base v original | 0.0104 | 1.1e-09 | 4.9e-08 |
| pearson_log1p v original | 0.011 | 2.7e-09 | 1.2e-07 |
| pearson_log v original | 0.00964 | 4.1e-09 | 1.8e-07 |
| icikt v pearson_base | 0.00819 | 3.8e-08 | 1.7e-06 |
| icikt v pearson_base_nozero | 0.00949 | 5.5e-08 | 2.5e-06 |
| icikt_complete v pearson_base_nozero | 0.00888 | 1.5e-06 | 6.8e-05 |
| icikt_complete v pearson_base | 0.00758 | 3.6e-06 | 1.6e-04 |
| pearson_base_nozero v kt_base | -0.00625 | 2.7e-05 | 1.2e-03 |
| pearson_base_nozero v icikt + pearson | -0.00629 | 6.1e-05 | 2.7e-03 |
| pearson_base_nozero v pearson_log1p | -0.00687 | 1.0e-04 | 4.7e-03 |
| pearson_base_nozero v icikt_complete + pearson | -0.00618 | 1.4e-04 | 6.5e-03 |
| pearson_base v icikt + pearson | -0.00499 | 1.6e-04 | 7.1e-03 |
| pearson_base_nozero v pearson_log | -0.00547 | 2.0e-04 | 9.0e-03 |
| pearson_base v kt_base | -0.00495 | 2.3e-04 | 1.0e-02 |
| pearson_base v original | 0.00547 | 2.6e-04 | 1.2e-02 |
| pearson_base v pearson_log1p | -0.00558 | 2.8e-04 | 1.3e-02 |
| pearson_base v icikt_complete + pearson | -0.00488 | 5.0e-04 | 2.2e-02 |
| icikt v icikt + pearson | 0.0032 | 5.1e-04 | 2.3e-02 |
| icikt_complete v icikt_complete + pearson | 0.0027 | 5.2e-04 | 2.3e-02 |
| pearson_base_nozero v original | 0.00417 | 1.4e-03 | 6.4e-02 |
| pearson_base v pearson_log | -0.00417 | 1.6e-03 | 7.0e-02 |
| icikt v icikt_complete + pearson | 0.00331 | 2.0e-03 | 9.1e-02 |
| icikt v pearson_log | 0.00402 | 4.5e-03 | 2.0e-01 |
| icikt v kt_base | 0.00324 | 7.4e-03 | 3.3e-01 |
| icikt_complete v pearson_log | 0.00341 | 1.5e-02 | 6.7e-01 |
| icikt_complete v kt_base | 0.00263 | 6.6e-02 | 1.0e+00 |
| icikt v pearson_log1p | 0.00262 | 3.3e-02 | 1.0e+00 |
| icikt_complete v icikt + pearson | 0.00259 | 2.6e-02 | 1.0e+00 |
| icikt_complete v pearson_log1p | 0.002 | 1.6e-01 | 1.0e+00 |
| pearson_log1p v pearson_log | 0.0014 | 3.5e-01 | 1.0e+00 |
| pearson_base v pearson_base_nozero | 0.0013 | 1.6e-01 | 1.0e+00 |
| pearson_log1p v icikt_complete + pearson | 0.000694 | 5.1e-01 | 1.0e+00 |
| pearson_log1p v kt_base | 0.000626 | 6.4e-01 | 1.0e+00 |
| icikt v icikt_complete | 0.000613 | 5.5e-01 | 1.0e+00 |
| pearson_log1p v icikt + pearson | 0.000585 | 4.7e-01 | 1.0e+00 |
| icikt + pearson v icikt_complete + pearson | 0.00011 | 8.8e-01 | 1.0e+00 |
| kt_base v icikt_complete + pearson | 6.83e-05 | 9.6e-01 | 1.0e+00 |
| kt_base v icikt + pearson | -4.14e-05 | 9.7e-01 | 1.0e+00 |
| pearson_log v icikt_complete + pearson | -0.000708 | 5.7e-01 | 1.0e+00 |
| pearson_log v kt_base | -0.000776 | 4.8e-01 | 1.0e+00 |
| pearson_log v icikt + pearson | -0.000818 | 5.2e-01 | 1.0e+00 |

### Feature-Feature Network Partitioning

**Figure S10.**
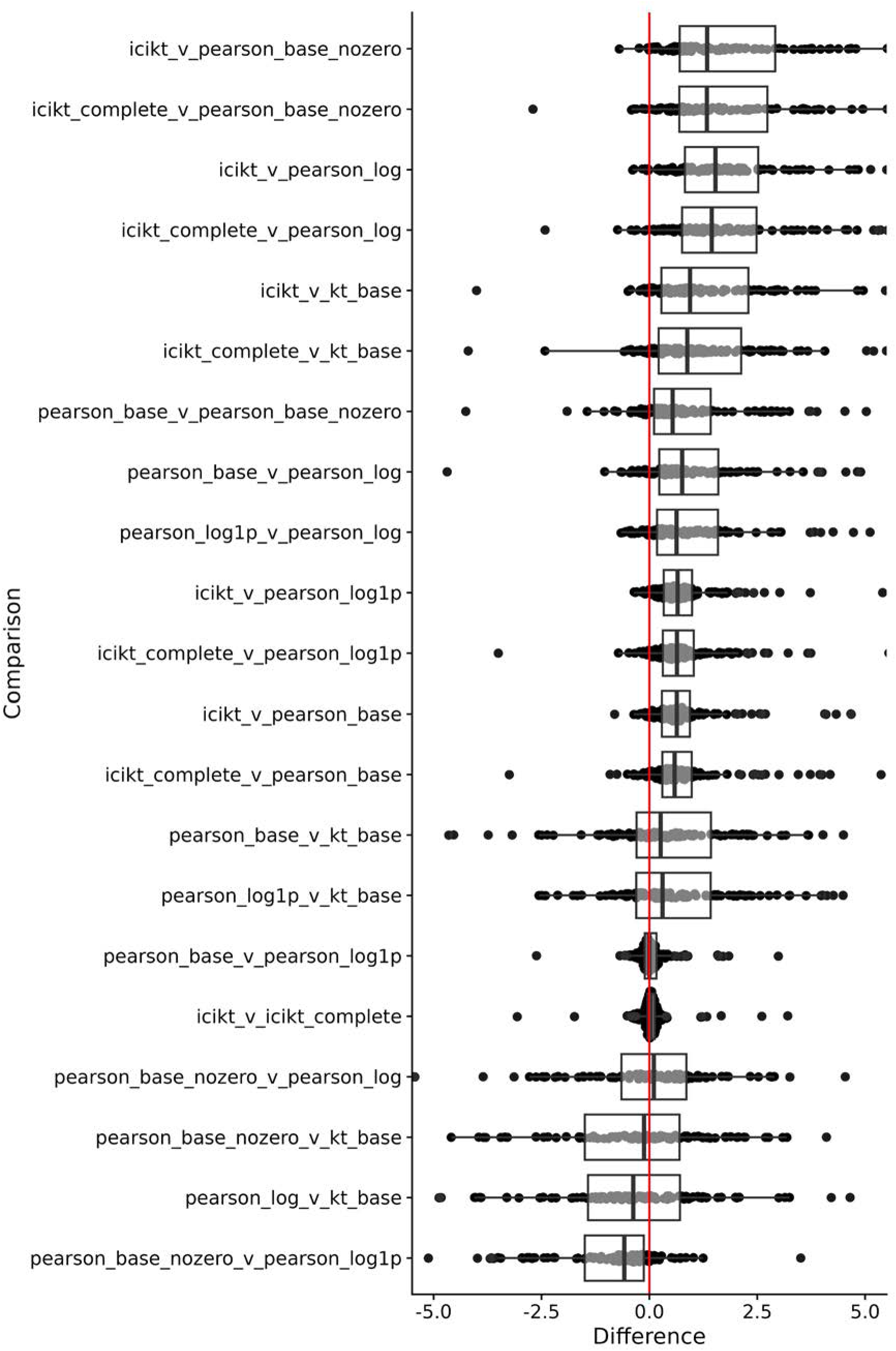
Boxplot and sina plots of paired differences of partitioning ratios across datasets. Red line indicates zero difference. Differences are calculated as *method*_1_ − *method*_2_.

**Table S2.** Paired t-test statistics comparing all the methods to each other. P-values were adjusted using the Bonferroni adjustment.

| Comparison | Difference | P-Value | P-Adjusted |
| --- | --- | --- | --- |
| icikt_complete v pearson_log | 2.3 | 2.7e-12 | 5.7e-11 |
| icikt v pearson_log | 2.35 | 3.9e-12 | 8.2e-11 |
| icikt v pearson_base_nozero | 2.51 | 4.1e-12 | 8.6e-11 |
| icikt_complete v pearson_base_nozero | 2.45 | 7.5e-12 | 1.6e-10 |
| icikt v kt_base | 1.7 | 1.2e-10 | 2.6e-09 |
| icikt_complete v kt_base | 1.63 | 3.7e-10 | 7.7e-09 |
| icikt v pearson_log1p | 1.15 | 8.6e-09 | 1.8e-07 |
| icikt_complete v pearson_log1p | 1.08 | 9.2e-09 | 1.9e-07 |
| icikt v pearson_base | 1.05 | 6.6e-08 | 1.4e-06 |
| icikt_complete v pearson_base | 0.976 | 6.8e-08 | 1.4e-06 |
| pearson_base v pearson_base_nozero | 1.41 | 7.7e-07 | 1.6e-05 |
| pearson_base_nozero v pearson_log1p | -1.33 | 1.7e-06 | 3.7e-05 |
| pearson_base v pearson_log | 1.31 | 3.1e-05 | 6.6e-04 |
| pearson_log1p v pearson_log | 1.18 | 3.4e-04 | 7.1e-03 |
| pearson_base v kt_base | 0.62 | 6.5e-04 | 1.4e-02 |
| pearson_log1p v kt_base | 0.534 | 1.0e-02 | 2.2e-01 |
| pearson_base_nozero v kt_base | -0.668 | 1.9e-02 | 4.0e-01 |
| pearson_log v kt_base | -0.676 | 4.4e-02 | 9.3e-01 |
| pearson_base v pearson_log1p | 0.0957 | 2.9e-01 | 1.0e+00 |
| icikt v icikt_complete | 0.0696 | 1.4e-01 | 1.0e+00 |
| pearson_base_nozero v pearson_log | -0.196 | 5.6e-01 | 1.0e+00 |

### Performance

**Figure S11.**
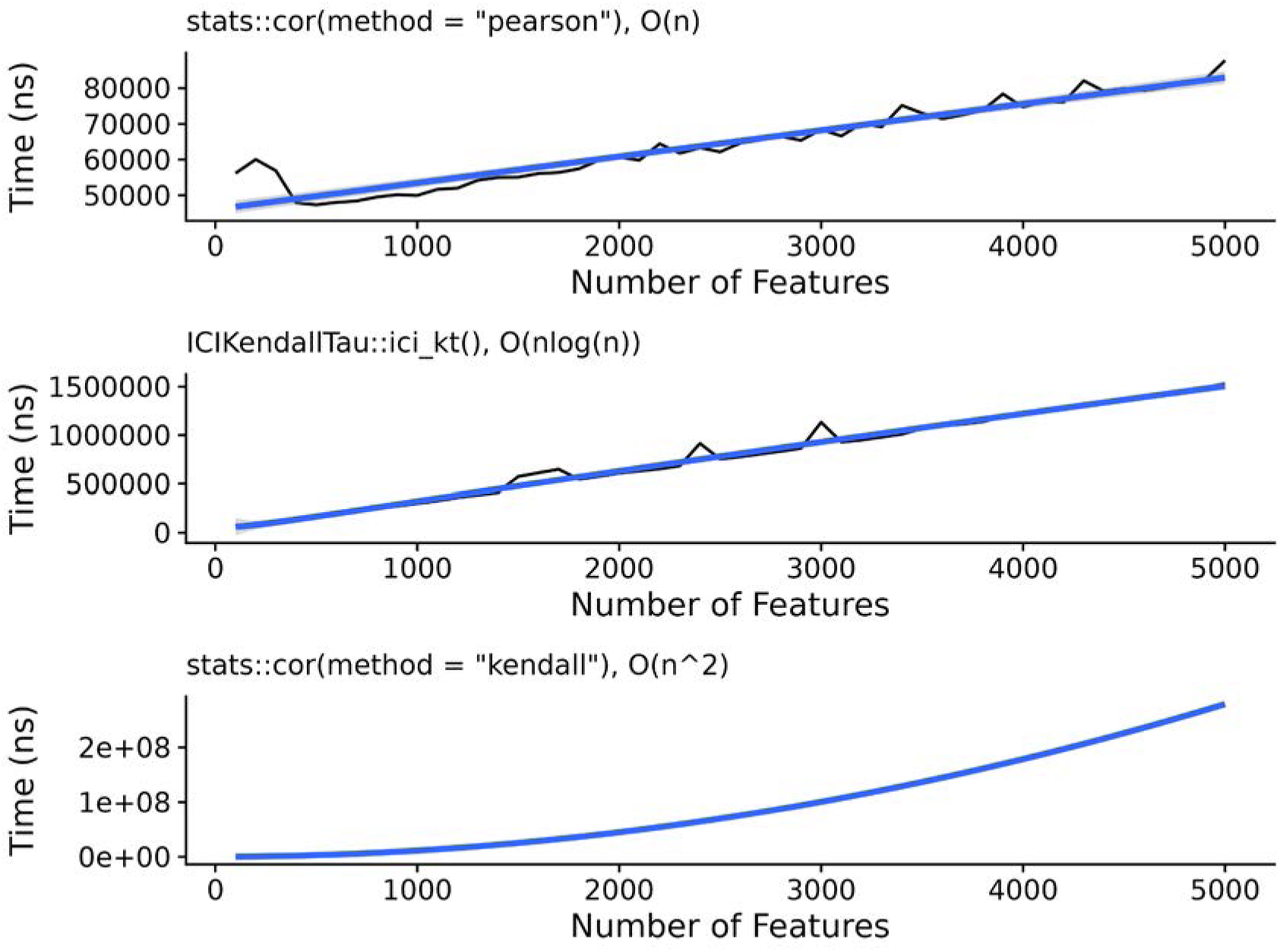
Time in seconds needed as a function of the number of features, with a fitted line for the assumed complexity for each of the methods tested, including R’s Pearson correlation, the ICI-Kt mergesort, and R’s Kendall-tau correlation algorithm.

